# Activating and repressing gene expression between chromosomes during stochastic fate specification

**DOI:** 10.1101/2022.07.11.499628

**Authors:** Elizabeth A. Urban, Chaim Chernoff, Kayla Viets Layng, Jeong Han, Caitlin Anderson, Daniel Konzman, Robert J. Johnston

## Abstract

DNA elements act across long genomic distances to regulate gene expression in processes including enhancer-promoter interactions and imprinting. During the gene-regulatory phenomenon of transvection in *Drosophila*, DNA elements on one allele of a gene act between chromosomes to increase or decrease expression of another allele of the gene. Despite the discovery of transvection over 60 years ago, little is known about its biological role. Furthermore, how different *cis* regulatory DNA elements contribute to the activation or repression of transvection at distinct times during development is unclear. Here, we studied the stochastic expression of *spineless* (*ss*) in developing photoreceptors in the fly eye to understand gene activation and repression between chromosomes. We identified a biological role for transvection in regulating expression of naturally occurring *ss* alleles. We characterized CRISPR-engineered deletions of sequences across the *ss* locus and identified DNA elements required for activating and repressing transvection. We found that different enhancers participated in transvection at different times during development to promote gene expression and specify cell fates. Bringing a silencer element on a heterologous chromosome into proximity with the *ss* locus “reconstituted” the gene, leading to repression. Our studies show that transvection regulates gene expression via distinct DNA elements at specific timepoints in development, with implications for genome organization and architecture.

## Introduction

Long-distance interactions between DNA regulatory elements are essential mechanisms for controlling gene expression. In mammals, enhancer-promoter interactions occur at distances as large as ∼1 Mb for genes including *Sonic Hedgehog*, *Hoxb*, and *Hoxd* (Ahn et al., 2014; Andrey et al., 2013; Lettice et al., 2003). In addition to long-range interactions along a single chromosome, DNA elements can act between chromosomes. In mice, olfactory receptor gene enhancers associate between chromosomes to form a super-enhancer that interacts with an active olfactory receptor gene (Monahan, 2019). Mammalian imprinting control regions act between chromosomes to ensure proper expression of maternal or paternal alleles (Herman et al., 2003; Sandhu et al., 2009). Aberrant interactions of DNA elements between chromosomes are observed in lymphoma and myeloma, where translocation of the *IgH* enhancer to chromosome 14 leads to activation of the *CCND1* locus on the homologous chromosome (Liu et al., 2008). However, little is known about the mechanisms controlling the action of regulatory DNA elements between chromosomes. Here we study interchromosomal gene regulation in *Drosophila melanogaster,* known as transvection, to understand its biological role and determine how noncoding DNA elements control gene expression between chromosomes at distinct times during development.

During transvection, the regulatory elements of a gene on one chromosome control expression of the same gene on a homologous chromosome. Transvection was originally discovered over 60 years ago by Ed Lewis at the *ubx* locus (Lewis, 1954) and has subsequently been described for a large number of *Drosophila* genes (Casares et al., 1997; Coulthard et al., 2005; Davison et al., 1985; Duncan, 2002; Gemkow et al., 1998; Hendrickson and Sakonju, 1995; Hopmann et al., 1995; Johnston and Desplan, 2014; Martinez-Laborda et al., 1992; Schoborg et al., 2013; Sipos et al., 1998; Southworth and Kennison, 2002; Tian et al., 2019). Transvection requires the colocalization, or pairing, of homologous chromosomes, which occurs in *Drosophila* in nearly all somatic cells throughout development (Stevens, 1906; Viets et al., 2019). Homologous chromosome pairing is driven by “button” elements, regions of high pairing affinity that are interspersed along chromosome arms (Fung et al., 1998; Gemkow *et al*., 1998; Hiraoka et al., 1993; Viets *et al*., 2019). With the exceptions of *Abd-b* and *spineless* (*ss*), pairing and transvection are typically disrupted by chromosomal rearrangements (Duncan, 2002; Gemkow *et al*., 1998; Hendrickson and Sakonju, 1995; Johnston and Desplan, 2014; Lewis, 1954; Sipos *et al*., 1998).

In most cases, transvection has been studied in the context of interactions between two paired mutant alleles. DNA elements on each allele act between chromosomes to rescue gene expression. Often, the enhancer of one mutant allele acts on the promoter of the other mutant allele to activate gene expression (“activating transvection”) (Duncan, 2002). An enhancer’s ability to act between chromosomes is often more efficient in the absence of a promoter in *cis* (Casares *et al*., 1997; Duncan, 2002; Gohl et al., 2008; Martinez-Laborda *et al*., 1992; Morris et al., 1999; Sipos *et al*., 1998). Insulator elements, which facilitate DNA looping interactions and block heterochromatin spreading (Phillips-Cremins and Corces, 2013), have also been linked to activating transvection (Fujioka et al., 2016; Kravchenko et al., 2005; Lim et al., 2018).

Transvection-related phenomena can also repress genes between chromosomes (“repressive transvection”), during processes including *brown* dominant silencing, *zeste-white* silencing, and pairing-sensitive silencing (Dreesen et al., 1991; Gans, 1953; Henikoff and Dreesen, 1989; Jack and Judd, 1979; Kassis, 2002; Zachar et al., 1985). DNA elements such as insulators and Polycomb Response Elements (PREs) have been linked to repressive transvection at many loci across the genome (Bantignies et al., 2003; Fauvarque and Dura, 1993; Fujioka et al., 1999; Gindhart and Kaufman, 1995; Kapoun and Kaufman, 1995; Kassis, 2002; Li et al., 2011; Muller et al., 1999; Shimell et al., 2000; Sigrist and Pirrotta, 1997). However, little work has focused on separating the functions of DNA elements that specifically contribute to activating and repressive transvection at a single locus. CRISPR provides the unique opportunity to precisely delete DNA elements from an endogenous gene locus and examine how they contribute to transvection.

As transvection has largely been studied in the context of mutant alleles, the biological role of transvection between naturally occurring alleles is unclear. It has been proposed that transvection ensures proper gene expression by enhancing the level of transcription between paired alleles (Goldsborough and Kornberg, 1996). However, a role for transvection between wild-derived alleles has yet to be experimentally demonstrated.

Regulation of *spineless* (*ss)* gene expression in the fly eye provides a paradigm to study transvection. In the fly retina, Ss is expressed stochastically in ∼67% of R7 photoreceptors (Ss^ON^ R7s) to activate expression of the photopigment Rhodopsin 4 (Rh4) and repress expression of Rhodopsin 3 (Rh3) (**Fig. 1A-B**). In the complementary 33% of R7s lacking Ss (Ss^OFF^ R7s), Rh3 is expressed (**Fig. 1A**) (Johnston and Desplan, 2014; Wernet et al., 2006). Therefore, Rh4 and Rh3 provide a readout for Ss^ON/OFF^ expression state in adult flies. Additionally, transvection between *ss* alleles can be measured quantitatively by assessing changes in the Rh4:Rh3 ratio.

**Figure 1.**
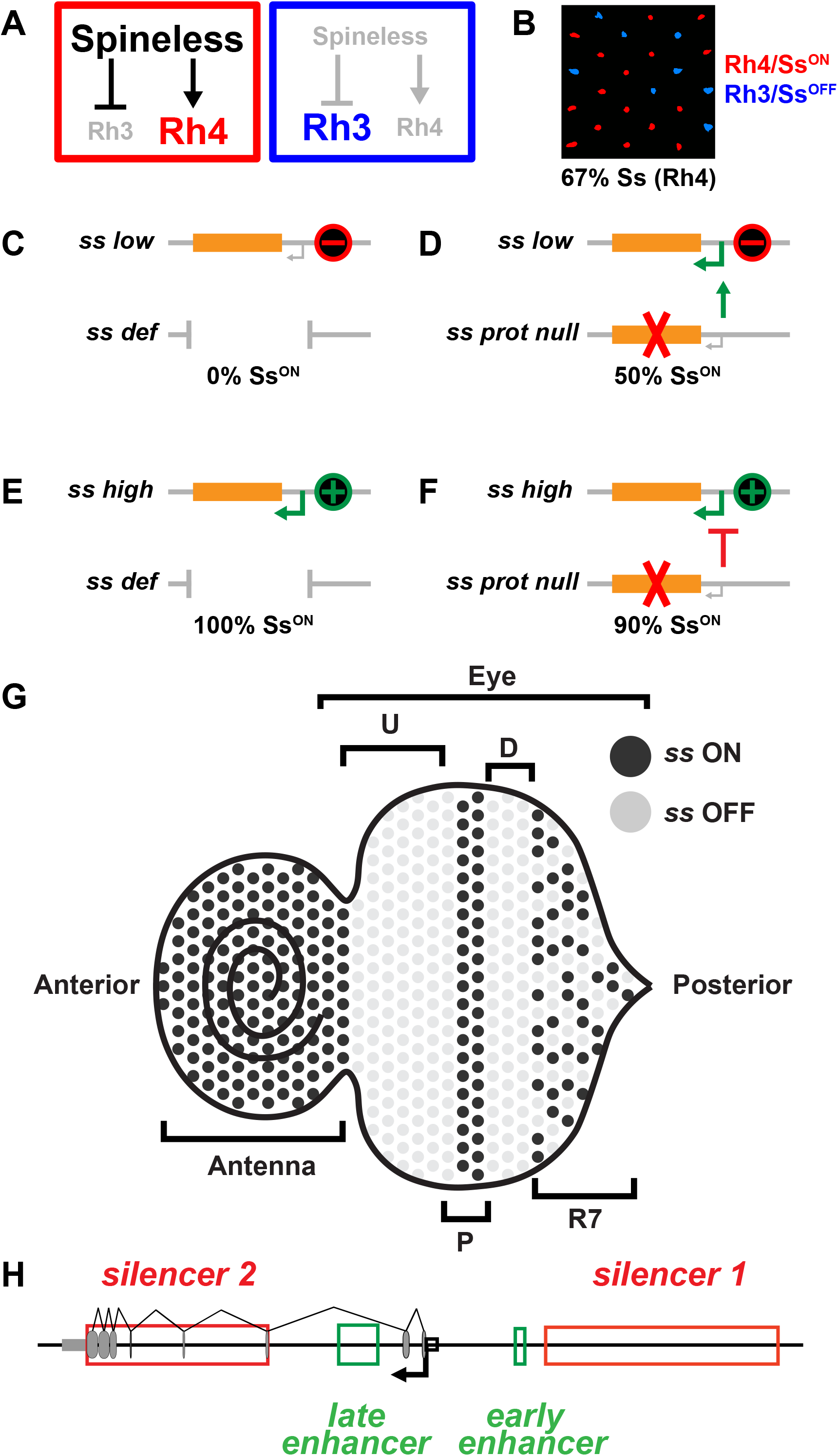
Transvection regulates *ss* expression during R7 subtype specification in the fly eye. **(A)** Spineless (Ss) activates Rh4 and represses Rh3. **(B)** Ss is expressed in ∼67% of R7s. Red: Rh4 (Ss^ON^ R7), blue: Rh3 (Ss^OFF^ R7). **(C-D)** Activating transvection: **(C)** *ss low*/ss def flies display 0% Ss^ON^ R7s. **(D)** *ss low/ss prot null* flies display 50% Ss^ON^ R7s, suggesting that the normal regulatory DNA elements from the *ss prot null* activate expression from the *ss low* allele. Red “X” indicates a mutation in exon 4 in *ss prot null* that yields a nonfunctional Ss protein product. Red “-” indicates a mutation in *ss low* that decreases *ss* expression. Green (active) and gray (inactive) arrows indicate the promoter and transcription start site. Orange box indicates *ss* protein coding region. **(E-F)** Repressive transvection: **(E)** *ss high/ss def* flies display 100% Ss^ON^ R7s. **(F)** *ss high/ss prot null* flies display 90% Ss^ON^ R7s, suggesting that the normal regulatory DNA elements from the *ss prot null* repress expression from the *ss high* allele. Red “X” indicates a mutation in exon 4 in *ss prot null* that yields a nonfunctional Ss protein product. Green “+” indicates a mutation in *ss high* that increases *ss* expression. Green (active) and gray (inactive) arrows indicate the promoter and transcription start site. Orange box indicates *ss* protein coding region. **(G)** 3^rd^ instar larval developing eye-antennal imaginal disc. Cells denoted by circles. *ss* RNA expressing cells in black, *ss*^OFF^ cells in grey. Eye disc development progresses from posterior to anterior. Cells proceed through a series of developmental stages: undifferentiated “U” (*ss*^OFF^), precursor “P” (*ss*^ON^), differentiating “D” (*ss*^OFF^), and terminal R7s “R7” (stochastic^ON/OFF^). Adapted from (Voortman, 2021). **(H)** Wildtype *spineless* gene locus. Red boxes = *silencer 1* and *silencer 2*, green boxes = *early* and *late enhancers*, grey ovals = exons, black arrow and box = promoter, grey rectangle = 3’UTR.

Mutations in *ss* regulatory DNA elements cause deviations from the wildtype 67% Ss^ON^ R7 ratio. Transvection activates and represses gene expression to determine a final Ss^ON^ R7 ratio (Johnston and Desplan, 2014). Flies with a hemizygous “*ss low”* mutant allele display *ss* expression in 0% of R7s (**Fig. 1C**). When a *ss low* allele is heterozygous with a *ss* protein coding null allele with normal regulatory regions, flies express Ss/Rh4 in 50% of R7s, showing that transvection activates *ss* expression (**Fig. 1D**)(Johnston and Desplan, 2014). Flies with a hemizygous “*ss high”* mutant allele display *ss* expression in 100% of R7s (**Fig. 1E**). When a *ss high* allele is heterozygous with the same *ss* protein coding null allele, flies express Ss/Rh4 in 90% of R7s, showing that transvection represses *ss* expression (**Fig. 1F**)(Johnston and Desplan, 2014).

Regulation of *ss* expression provides a unique opportunity to understand when and where transvection occurs during development. We found that expression of *ss* in a random subset of R7s is controlled by a dynamic interplay between transcription and chromatin earlier in development (Voortman, 2021). Photoreceptor specification occurs through a wave of differentiation during larval development (Ready et al., 1976; Tomlinson and Ready, 1987a; b; Wolff and Ready, 1991). Due to this wave-like nature, multiple stages of eye development can be observed at a single timepoint. *ss* is regulated in four stages during development (**Fig. 1G**). Initially, *ss* is off in undifferentiated cells (**Fig. 1G, “U”)**. An *early enhancer* activates *ss* transcription in all precursor cells to open the gene locus (**Fig. 1G, 1H, “P”)**. *ss* turns off in differentiating cells (**Fig. 1G, “D”)**. Finally, a *late enhancer* reactivates *ss* expression in a subset of R7s dependent on chromatin state (**Fig. 1G, 1H, “R7”**). Ss^ON^ R7s express *ss* and have open chromatin at the *ss* locus, whereas Ss^OFF^ R7s repress *ss* and have compact chromatin (Voortman, 2021). The regulation of *ss* by two enhancers that function at distinct timepoints provides a model to elucidate the spatiotemporal roles of transvection.

Here, we studied transvection at the *ss* locus to understand activation and repression between chromosomes. We identified a biological role for transvection in controlling the expression of naturally occurring *ss* alleles in the fly retina. Using precise deletions of DNA regions across the *ss* locus, we identified elements that mediate transvection between *ss* alleles and separated the activating and repressing components of transvection. Activation of *ss* between chromosomes required an intact enhancer and promoter on the same chromosome, while repression of *ss* between chromosomes required two PREs and an insulator on both chromosomes. We separated the spatiotemporal properties of *ss* transvection, finding that transvection occurs early in precursors via the *early enhancer* and late during terminal R7 subtype specification via the *late enhancer*. Finally, we observed *ss* pairing and transvection in the presence of substantial chromosomal rearrangements with breakpoints within the *ss* locus, showing that physical proximity can “reconstitute” the *ss* gene between chromosomes to promote normal expression.

## Results

### Transvection between wild-derived alleles yields an intermediate proportion of SsON R7s

The role of transvection in regulating the expression of naturally occurring wild-derived alleles is poorly understood. Wild-derived alleles of *ss* from the Drosophila Genome Resource Panel (DGRP) display wide variation in their proportions of Ss^ON^ R7s (Anderson et al., 2017; Mackay et al., 2012). To limit genetic variability arising from other chromosomes, we isogenized the third chromosome containing *ss*. Flies that were homozygous for wild-derived *ss* alleles displayed Ss^ON^ R7 ratios ranging from 41% to 74% (**Fig. 2A-D**). To investigate how wild-derived *ss* alleles interact via transvection, we crossed parent flies and examined Ss^ON^ R7 ratios in heterozygous progeny. We hypothesized that, if the two alleles did not perform transvection, their heterozygous progeny would exhibit either (1) the same proportion of Ss^ON^ R7s as the higher allele, or (2) a higher proportion of Ss^ON^ R7s than either of the parents due to independent expression of each parental allele. Alternatively, if the two alleles activated and repressed each other via transvection, their heterozygous progeny would exhibit an intermediate proportion of Ss^ON^ R7s.

**Figure 2.**
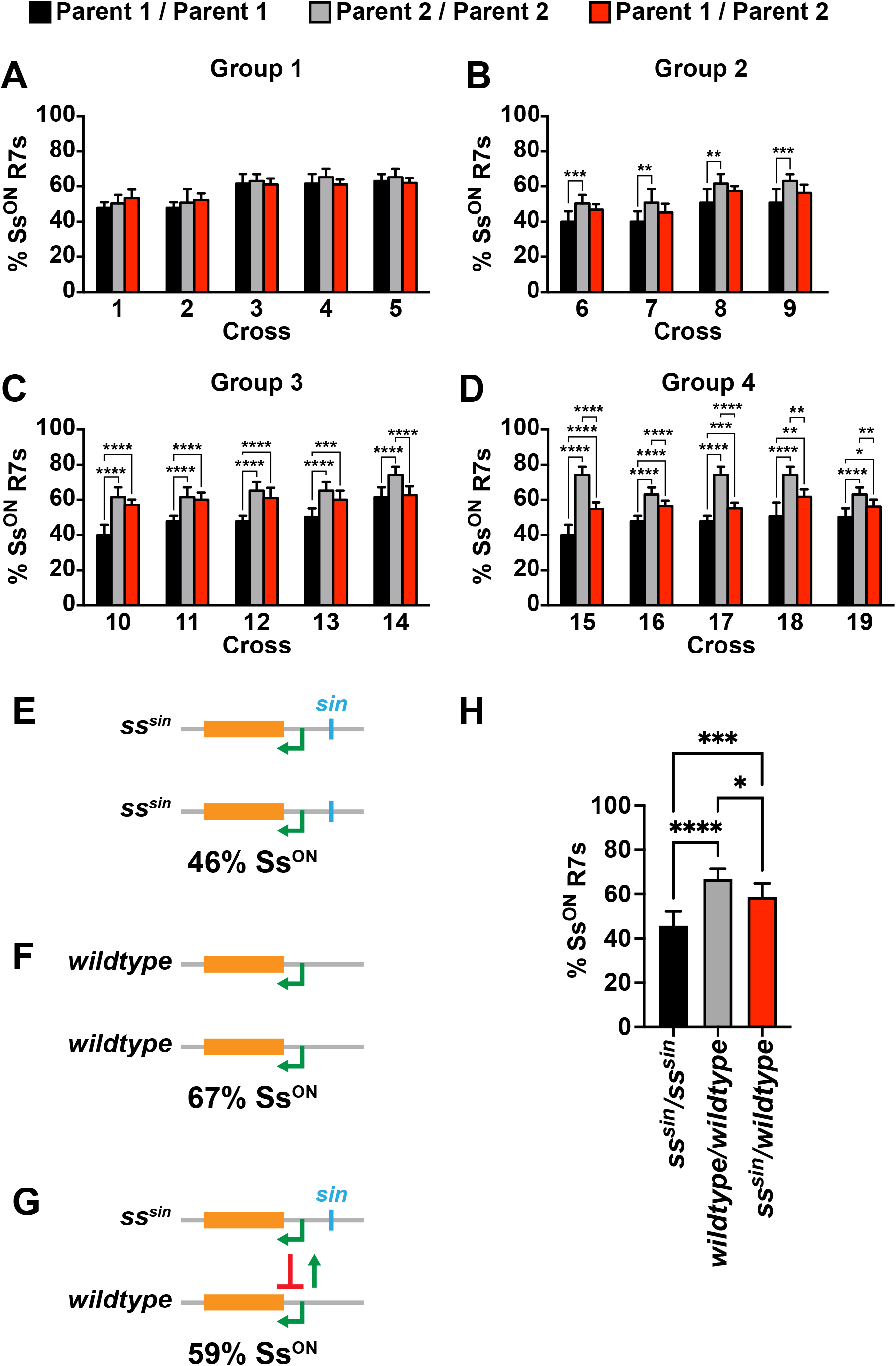
Transvection between wild-derived alleles yields an intermediate proportion of Ss^ON^ R7s. **(A-D)** Graphs of % Ss^ON^ R7s for; homozygous parent 1 (black), homozygous parent 2 (grey), and progeny of parent 1 / parent 2 (red) for Groups 1-4. ****=p<0.0001, ***=p<0.001, **=p<0.005, *=p<0.05, unpaired t-test. n = 7-10 retinas/genotype. **(A)** Group 1: % Ss^ON^ R7s of parents are not statistically different from each other; progeny are not statistically different from either parent. **(B)** Group 2: % Ss^ON^ R7s of parents are statistically different from each other; % Ss^ON^ R7s of progeny are not statistically different from either parent. **(C)** Group 3: % Ss^ON^ R7s of parents are statistically different from each other; % Ss^ON^ R7s of progeny are statistically different from one parent. **(D)** Group 4: % Ss^ON^ R7s of parents are statistically different from each other; % Ss^ON^ R7s of progeny are statistically different from both parents. **(E-G) (G)** *ss^sin^*/*wildtype* flies display intermediate % Ss^ON^ R7s compared to **(E)** *ss^sin^*/*ss^sin^* and **(F)** *wildtype/wildtype* flies. Green arrow indicates transcription start site. Orange box indicates *ss* protein coding region. **(H)** Quantification of **E-G.** ****=p<0.0001, ***=p<0.001, **=p<0.005, *=p<0.05, one-way ANOVA. n = 8- 11 retinas/genotype.

We determined the proportion of Ss^ON^ R7s for 19 heterozygous conditions and classified phenotypes into four categories. In Group 1, the proportion of Ss^ON^ R7s of both parents and their progeny were not statistically different from each other (**Fig. 2A**). In Groups 2-4, the parents had statistically different proportions of Ss^ON^ R7s (**Fig. 2B-D**). The proportions of Ss^ON^ R7s in Group 2 progeny were not statistically different from either parent (**Fig. 2B**), the proportions of Ss^ON^ R7s of Group 3 progeny were statistically different from one parent (**Fig. 2C**), and the proportions of Ss^ON^ R7s of Group 4 progeny were statistically different from both parents (**Fig. 2D**). In all cases, heterozygous progeny had a proportion of Ss^ON^ R7s that fell between the ratios of the parents or were not significantly different from the ratios of the parents (**Fig. 2A-D**). Thus, flies that are heterozygous for wild-derived *ss* alleles exhibit an intermediate proportion of Ss^ON^ R7s, suggesting that naturally occurring *ss* alleles regulate each other via transvection.

To further investigate transvection between wild-derived alleles, we examined a *ss* allele in which we used CRISPR to insert a natural variant (*sin*) into a laboratory stock (Anderson et al., 2017). Flies that are homozygous for *sin* displayed 46% Ss^ON^ R7s (**Fig. 2E, 2H**), while wildtype flies lacking *sin* had 67% Ss^ON^ R7s (**Fig. 2F, 2H**) (Anderson *et al*., 2017). *sin/wildtype* heterozygotes had 59% Ss^ON^ R7s (**Fig. 2G-H**), intermediate between the parental alleles.

Together, transvection regulates *ss* from naturally occurring alleles to yield intermediate proportions of Ss^ON^ R7s, suggesting a biological role for transvection in patterning the *Drosophila* retina.

### Two enhancers promote *ss* expression via transvection at two developmental timepoints

The natural variant, *sin,* affects the *early enhancer,* one of two enhancers that are required for stochastic expression of *ss.* The *early enhancer* activates *ss* expression in precursors, whereas the *late enhancer* activates expression in terminal R7s (**Fig. 1G-H**). We examined flies with mutations in these enhancers to understand temporal control of *ss* expression via transvection during development.

To test the roles of the *early* and *late enhancers* in transvection, we tested alleles with deletions of the *early enhancer* (*early enhΔ*) or the *late enhancer* (*late enhΔ*)(Voortman, 2021). For these experiments, we used RNA fluorescent *in situ* hybridization (FISH) to assess changes in early *ss* expression in precursor cells in 3^rd^ instar larvae. We designed single stranded probes against a region of the transcript deleted in the *prot null* allele (i.e. part of intron 3, all of exon 4, and part of intron 4). These probes detected nascent *ss* transcripts from the *wildtype, early enhΔ,* and *late enhΔ* alleles, but not the *prot null* allele (**Fig. S1A**). To quantify changes in *ss* expression in precursor cells, we measured the density of nascent *ss* RNA spots as a proxy for the number of active *ss*^ON^ precursor cells. We measured the density of *ss*^ON^ precursors in a sliding window across the length of the disc to account for any stretching or compression due to mounting (**Fig. S1D**). To quantify *ss* expression in mature R7s, we assessed Rh3 and Rh4 expression in adult retinas, as they faithfully report Ss expression in R7s (i.e. Ss^ON^ = Rh4; Ss^OFF^ = Rh3) (**Fig. 1B**).

We assessed expression from the *prot null* allele, which is a deletion of exon 4 and parts of introns 3 and 4, leading to an inactive protein. As the *prot null* allele has intact regulatory regions, we expected to observe RNA expression from the *prot null* allele that resembled *wildtype*. Using probes that targeted the entire *ss* transcript, we detected nascent *ss* RNA expression in precursors in *prot null/def* flies (**Fig S1A, S1C’; *ss* RNA (full)**), showing that this allele retains intact *cis-*regulatory elements that drive normal RNA expression. When using probes that targeted the deleted region of the *prot null* allele, we did not observe RNA expression signal in precursors in *prot null/def* flies (**Fig S1A, S1C”; *ss* RNA (subset)**), consistent with the absence of this region in the *prot null* allele. We observed no Rh4-expressing R7s in *prot null/def* adult animals, consistent with the loss of protein function (**Fig. S1C’’’**). *wildtype/ss def* and *wildtype/prot null* flies displayed similar *ss* expression in precursors and R7s, consistent with the *prot null* having normal regulatory regions (**Fig. 3A, 3G**). These results suggest that (1) the *prot null* allele contains functional *cis-*regulatory elements that drive expression at the correct time and place, and could alter expression from other alleles, (2) our *ss* RNA (subset) probes do not detect expression in precursors from the *prot null* allele, and thus any RNA observed in heterozygous flies must arise from the non-*prot null* allele, and (3) the *prot null* allele does not produce Ss protein that drives Rh4 expression in Ss^ON^ R7s, and thus any Rh4/Ss^ON^ R7s observed in heterozygous flies indicate expression from the non-*prot null* allele.

**Figure 3.**
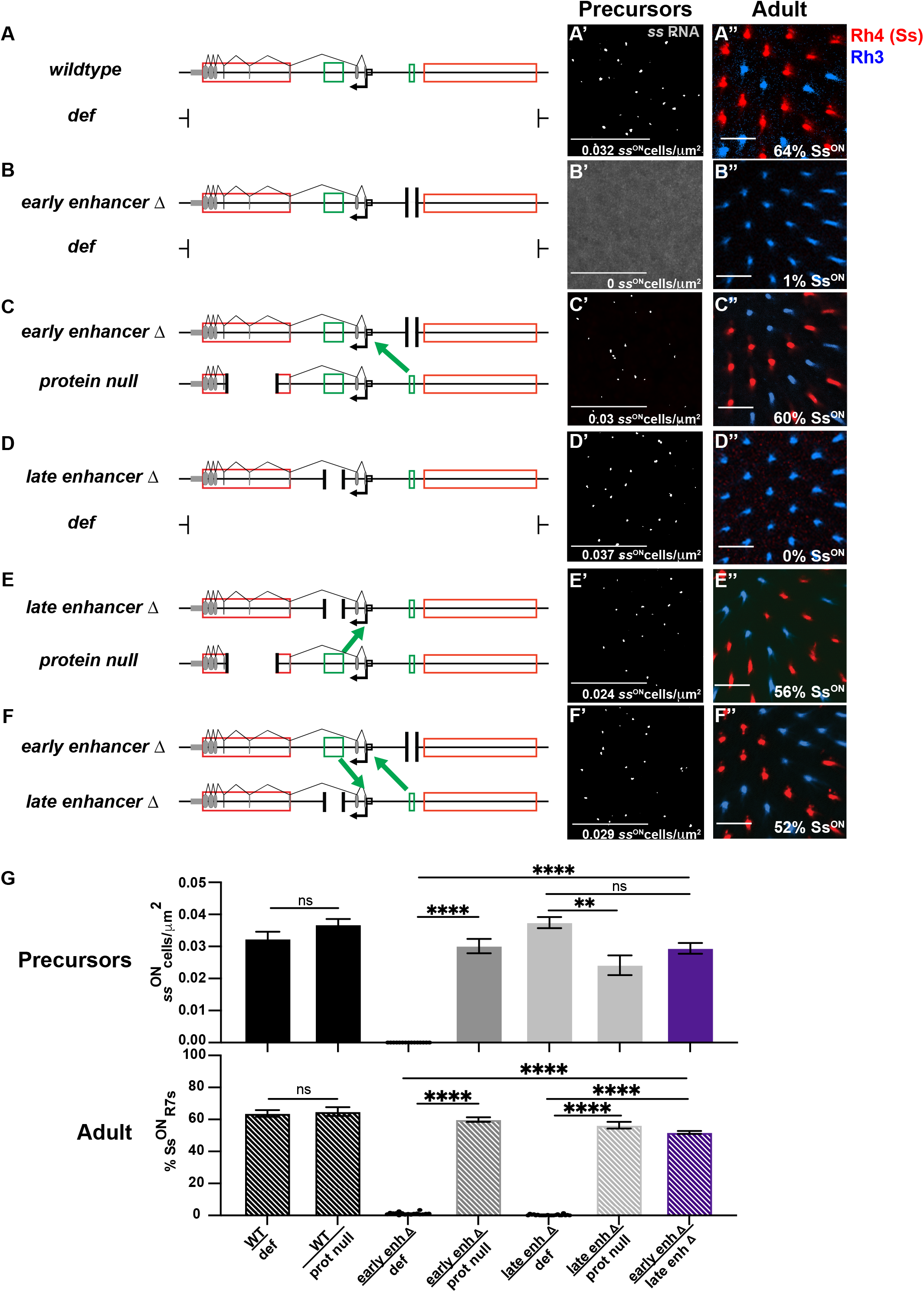
Two enhancers activate transvection at distinct times during development. **(A-F).** Schematics for **(A)** *wildtype/def,* **(B)** *early enhancer Δ/def,* **(C)** *early enhancer Δ/protein null*, **(D)** *late enhancer Δ/def,* **(E)** *late enhancer Δ/protein null,* and **(F)** *early enhancer Δ/late enhancer Δ*. Green arrows indicate activating transvection is occurring between alleles from the early or late enhancer. Gray ovals: exons, black arrow and box: promoter, red boxes: silencers, green boxes: enhancers, grey box: 3’UTR. **(A’-F’)** Representative images of RNA FISH using subset probes (**Fig. S1**) in precursor R7s. Grey dots represent nascent transcript expression. Due to homologous chromosome pairing, each cell expresses only 1 nascent site per nuclei. **(A’)** *wildtype/def,* **(B’)** *early enhancer Δ/def,* **(C’)** *early enhancer Δ/protein null*, **(D’)** *late enhancer Δ/def,* **(E’)** *late enhancer Δ/protein null,* and **(F’)** *early enhancer Δ/late enhancer Δ*. Scale bars:15 μm **(A”-F”)** Representative images and % Ss^ON^ R7s (Rh4) in adult retinas. Antibody staining against Rh4(Ss): red and Rh3: blue. **(A”)** *wildtype/def,* **(B”)** *early enhancer Δ/def,* **(C”)** *early enhancer Δ/protein null*, **(D”)** *late enhancer Δ / def,* **(E”)** *late enhancer Δ/protein null,* and **(F”)** *early enhancer Δ/late enhancer Δ*. Scale bars:15 μm. **(G)** Quantification of *ss* expression in precursors and adult R7s from figure **A-F**. Error bars: SEM. P- values calculated by one-way ANOVA with Tukey’s multiple comparisons. ****=p<0.0001, ***=p<0.001, **=p<0.005, *=p<0.05.

We next addressed the roles of the *early enhancer* and *late enhancer* in transvection. We predicted that, if transvection occurs early in precursor cells, we would see a change in the number of cells expressing *ss* RNA. Change in expression early is linked to a change in late Ss^ON/OFF^ (Voortman, 2021). For example, a decrease in the number of precursor cells expressing *ss* will be reflected as a decrease in terminal Ss^ON^ R7s. However, if transvection occurs late in development, we expected to observe no change in expression in precursor cells, but a change in the terminal % Ss^ON^ R7s. We previously found that all precursors express *ss* in homozygous wildtype flies (Voortman, 2021).

The *early enhancer* drives early *ss* expression in precursors, which enables the *late enhancer* to drive expression in terminal Ss^ON^ R7s (Voortman, 2021). *early enhΔ/def* flies displayed no expression in precursors and 1% Ss^ON^ R7s (**Fig. 3B, G**). *early enhΔ/prot null* flies displayed an increase in *ss* expression in precursors and % Ss^ON^ R7s compared to *early enhΔ/def* (**Fig. 3C, G**), suggesting that the *early enhancer* can drive transvection early in development, enabling late *ss* expression in Ss^ON^ R7s.

The *late enhancer* is not required for early *ss* expression in precursors but is required for late *ss* expression in Ss^ON^ R7s (Voortman, 2021). *late enhΔ/def* flies displayed *ss* expression in precursors, but 0% Ss^ON^ R7s (**Fig. 3D, G**). *late enhΔ/prot null* flies displayed *ss* expression in precursors and an increase in Ss^ON^ R7s (56%; **Fig. 3E, G**), suggesting that the *late enhancer* activates expression via transvection late to promote Ss^ON^ R7 fate.

To test whether the *early enhancer* and *late enhancer* could reciprocally promote expression, we examined *early enhΔ/late enhΔ* flies. Whereas *early enhΔ/def* flies did not express *ss* in precursors or R7s (**Fig. 3B**) and *late enhΔ/def* flies expressed *ss* in precursors but not R7s (**Fig. 3D**), *early enhΔ/late enhΔ* flies expressed *ss* in precursors and 52% Ss^ON^ R7s (**Fig. 3F, G**), consistent with transvection from the *early enhancer* and *late enhancer* to promote expression at both developmental timepoints.

Together, these data suggest that the *early* and *late enhancers* participate in transvection at distinct times during development. The *early enhancer* activates expression between chromosomes in precursors, while the *late enhancer* activates expression between chromosomes in terminal R7s to specify Ss^ON^ R7 fate.

### Enhancers and the promoter on the same chromosome regulate activating transvection

In many cases of transvection, enhancers activate expression between chromosomes more efficiently when an enhancer does not have a promoter on the same chromosome (Bateman et al., 2012a; Blick et interactions of the enhancers and the promoter in activating transvection at the *ss* locus, we assessed the effects of a *promoter* deletion allele that deletes 2 kb encompassing the *ss* transcriptional start (Voortman, 2021). *promoterΔ/def* flies displayed no *ss* expression in precursors and 1% Ss^ON^ R7s (**Fig. 4A**), suggesting a near complete ablation of promoter function. The minimal expression in adults may be due to cryptic promoter activity.

**Figure 4.**
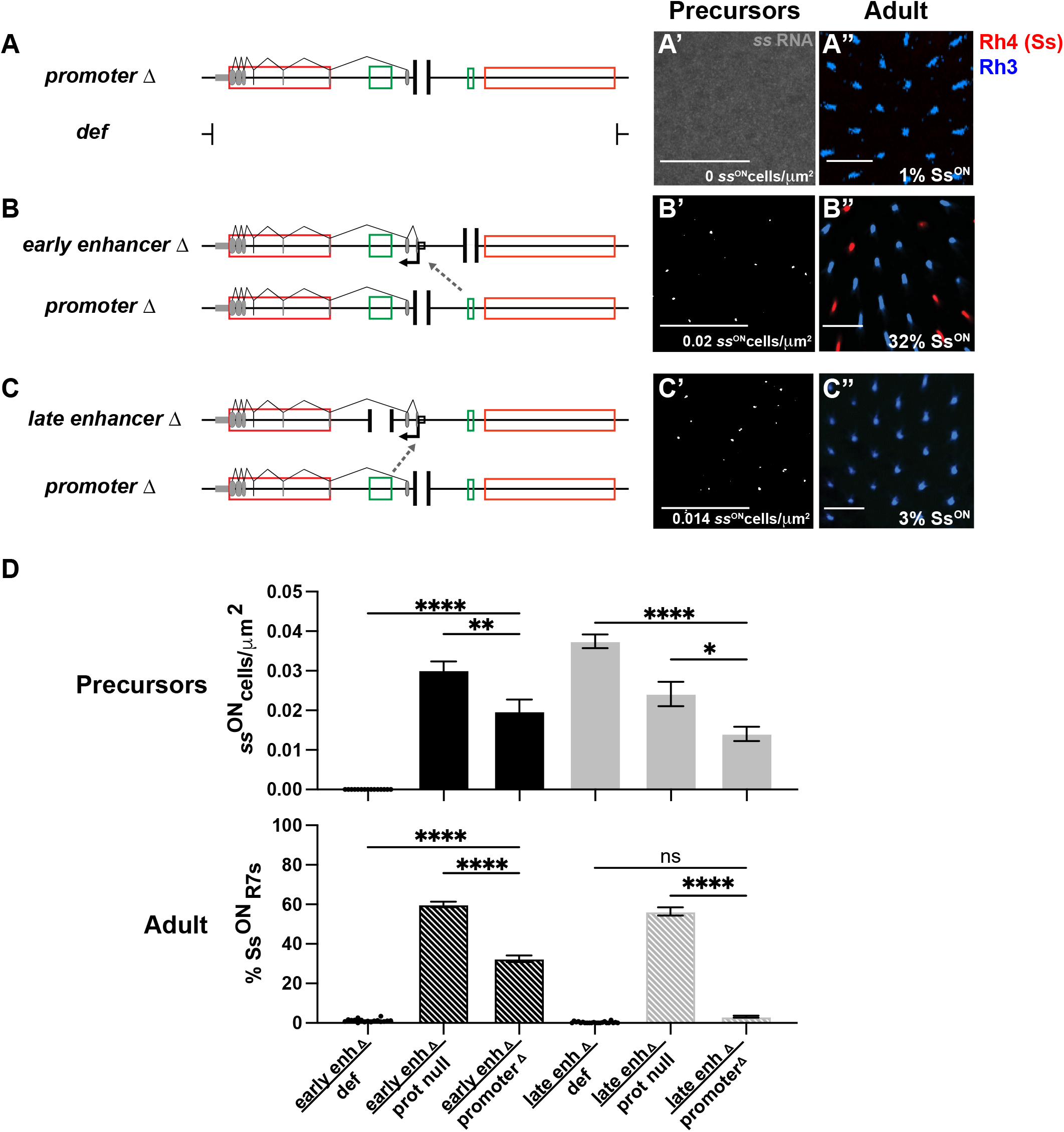
Activating transvection requires an enhancer and promoter on the same chromosome. **(A-C)** Schematics for (A) *promoter Δ/def,* (B) *early enhancer Δ/promoter Δ*, and (C) *late enhancer Δ/promoter Δ.* Green arrows indicate activating transvection is occurring between alleles. Grey dashed arrow indicates failure to perform transvection. **(A’-C’)** Representative images of RNA FISH using subset probes in precursor cells. Grey dots are nascent transcript expression. Each cell expresses only 1 nascent site (nuclei not shown). **(A’)** *promoter Δ/def,* **(B’)** *early enhancer Δ/promoter Δ*, **(C’)** *late enhancer Δ/promoter Δ.* Scale bar: 15 μm. **(A”-C”)** Representative images and % Ss^ON^ R7s (Rh4). Antibody staining against Rh4(Ss):red and Rh3:blue. **(A”)** *promoter Δ/def,* **(B”)** *early enhancer Δ/promoter Δ*, **(C”)** *late enhancer Δ/promoter Δ.* Scale bar: 15 μm. **(D)** Quantification of RNA FISH and % Ss^ON^ R7s from **A-C**. Error bars: SEM. P-values calculated one- way ANOVA with Tukey’s multiple comparisons. ****=p<0.0001, ***=p<0.001, **=p<0.005, *=p<0.05.

*early enhΔ/def* flies did not express *ss* in precursors or R7s (**Fig. 4D**). *early enhΔ/prot null* flies displayed a robust increase in *ss* expression in precursors and % Ss^ON^ R7s (**Fig. 4D**). *early enhΔ/promoterΔ* flies promoted expression in precursors, but in significantly less cells, and displayed only 32% Ss^ON^ R7s (**Fig. 4B, D**), suggesting that the *early enhancer* requires the *promoter* to efficiently promote activating transvection in precursors and subsequent specification of Ss^ON^ R7s

*late enhΔ/def* flies displayed *ss* expression in precursors and a complete loss of Ss^ON^ R7s (**Fig. 4D**). *late enhΔ/prot null* flies displayed *ss* expression in precursors and an increase in % Ss^ON^ R7s (**Fig. 4D**). *late enhΔ/promoterΔ* flies displayed reduced expression in precursors that may be due to repressive transvection or genetic background effects (**Fig. 4C, D**). *late enhΔ/promoterΔ* flies displayed almost no Ss^ON^ R7s (Ss^ON^ = 3%; **Fig. 4C, D**), suggesting that the *late enhancer* requires the *promoter* to efficiently promote activating transvection in Ss^ON^ R7s.

Together, these data suggest that the promoter together with the enhancers on the same chromosome promote activating transvection at the *ss* locus.

### Repression of *ss* between chromosomes requires distinct elements within a silencer

To investigate the roles of regulatory DNA elements in repressive transvection between *ss* alleles, we examined *silencer 1* at the 5’ end of the *ss* locus. *silencer 1* contains an insulator element (Viets *et al*., 2019) and two Polycomb Response Elements (PREs) (Voortman, 2021) (**Fig. 5A**). Insulators are *cis*-regulatory DNA elements that are bound by insulator proteins. Insulators play a role in DNA looping to block heterochromatin spreading or limit enhancer-promoter interactions (Gurudatta and Corces, 2009). PREs are bound by members of the repressive Polycomb Group Complex and are sites of heterochromatin nucleation (Czermin et al., 2002; Duncan, 1998; Geisler and Paro, 2015; Lewis, 1978; Piunti and Shilatifard, 2016; Schwartz and Pirrotta, 2013; Simon and Kingston, 2013). Deletions of both *PREs* or the entire *silencer 1* lead to increases in the proportion of Ss^ON^ R7s (Voortman, 2021).

**Figure 5.**
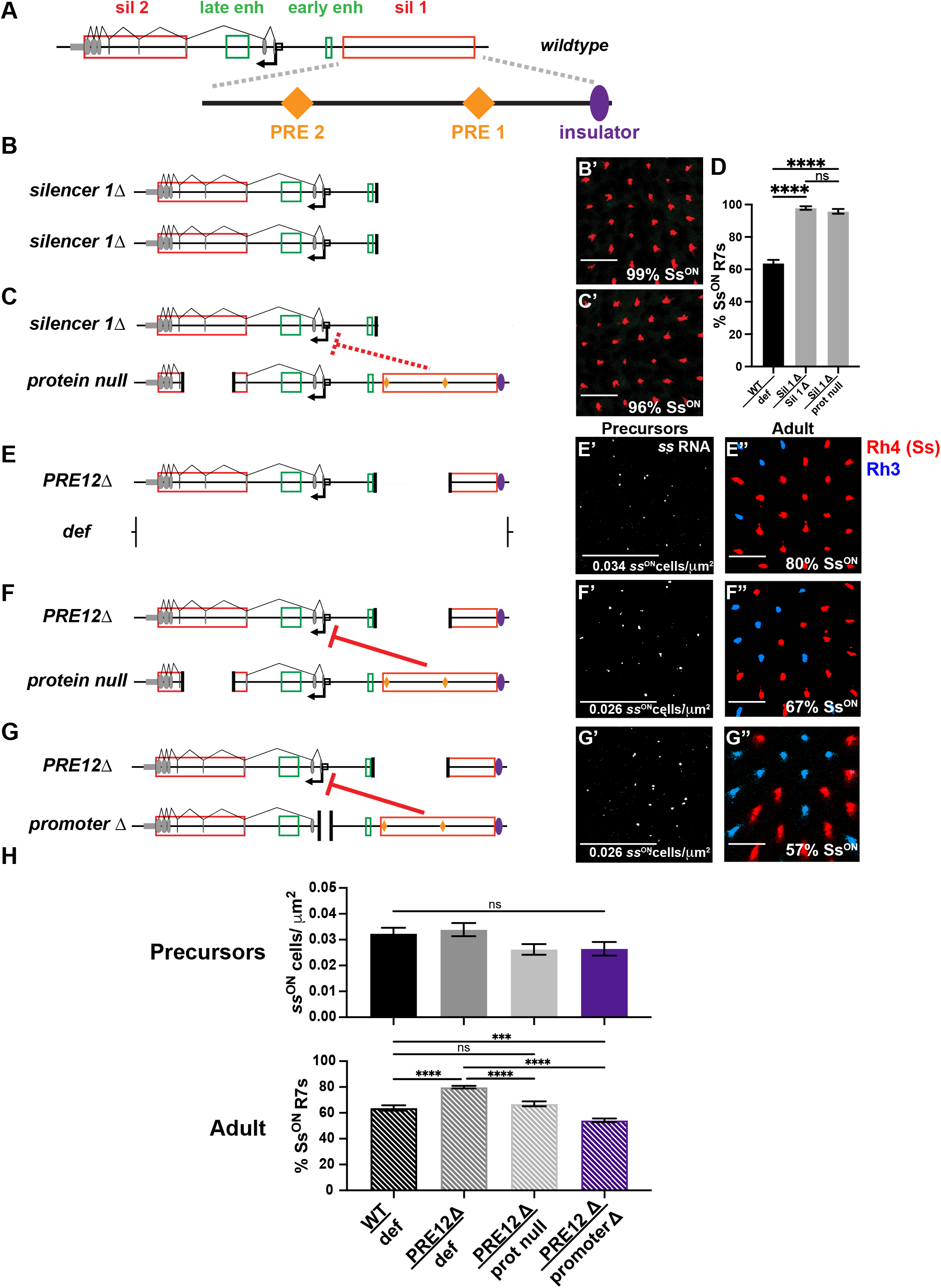
Repressive *ss* transvection requires two PREs and an insulator element. **(A)** Schematic of the wildtype *ss* locus with an expanded view of *silencer 1*, which contains two PREs (orange diamonds) and an insulator (purple oval). sil 1: *silencer1*, early enh: *early enhancer*, late enh: *late enhancer*, sil 2: *silencer2*. Gray ovals: exons, black arrow and square: promoter. **(B-F)** Purple oval: insulator, orange diamonds: PREs. Red bar indicates that *silencer 1* is performing repressing transvection. Dotted red bar indicates that *silencer 1* on *protein null* is not performing repressing transvection on *silencer 1Δ*. **(B-C)** Schematics and representative images of % Ss^ON^ R7s in adult retinas. Antibody staining against Rh4(Ss):red and Rh3:blue for **(B’)** *silencer1Δ/silencer1Δ* and **(C’)** *silencer 1Δ/protein null.* Scale bars: 15 μm. **(D)** Quantification of % Ss^ON^ R7s for **B-C**. P values calculated by one-way ANOVA with Tukey’s multiple comparison test. ****=p<0.0001, ***=p<0.001, **=p<0.005, *=p<0.05. Error bars: SEM. **(E-G)** Schematics for **(E)** *PRE12Δ/ss def*, **(F)** *PRE12Δ/protein null,* and **(G)** *PRE12Δ/promoterΔ*. **(E’-G’)** Representative images of RNA FISH using subset probes in precursor cells. Grey dots are sites of nascent transcript expression. Each cell expresses only 1 nascent site (nuclei not shown) for **(E’)** *PRE12Δ/ss def*, **(F’)** *PRE12Δ/protein null,* and **(G’)** *PRE12Δ/promoterΔ .* Scale bars: 15 μm. **(E”-G”)** Representative images of % Ss^ON^ R7s in adult retinas. Antibody staining against Rh4(Ss):red and Rh3:blue for **(E”)** *PRE12Δ/ss def*, **(F”)** *PRE12Δ/protein null,* and **(G”)** *PRE12Δ/promoterΔ .* Scale bars: 15 μm. **(H)** Quantification of RNA FISH and % Ss^ON^ R7s from **E-G**. Error bars: SEM. P-values calculated one-way ANOVA with Tukeys multiple comparisons. ****=p<0.0001, ***=p<0.001, **=p<0.005, *=p<0.05.

We examined the effects of a 36 kb deletion of *silencer 1* (*silencer 1Δ*) (Thanawala et al., 2013) on transvection at the *ss* locus. *silencer 1Δ* homozygotes displayed 99% Ss^ON^ R7s (**Fig. 5B, 5D**)(Johnston and Desplan, 2014). *silencer 1Δ/prot null* flies similarly had 96% Ss^ON^ R7s (**Fig. 5C, D**), suggesting that repressive transvection does not occur when all of the elements of *silencer1* are deleted (Johnston and Desplan, 2014).

We next examined a 17 kb deletion that removed both PRE1 and PRE2 but left *insulator1* intact (*PRE12Δ*). *PRE12Δ*/*ss def* flies displayed no change in early *ss* expression in precursors, but an increase to 80% Ss^ON^ R7s (**Fig. 5E, 5H**)(Viets *et al*., 2019). *PRE12Δ/prot null* flies displayed no change in *ss* expression in precursors, but a decrease in the proportion of Ss^ON^ R7s to 67% (**Fig. 5F, H**). As *insulator1* is present in *PRE12Δ* but absent in *silencer 1Δ,* these data suggest that *insulator1* facilitates repressive transvection.

The promoter together with the enhancer on the same chromosome promote *ss*-activating transvection. We tested the role of the promoter in repressive transvection. *PRE12Δ/promoterΔ* flies displayed no change in *ss* expression in precursors, but a decrease in the proportion of Ss^ON^ R7s to 54% (**Fig. 5G, H**), indicating that *silencer1* does not require the *ss promoter* on the same chromosome for repressive transvection and that repressive transvection at the *ss* locus occurs more efficiently in the absence of a promoter (**Fig. 5H**). Together, these data suggest that activating and repressing transvection are mediated by different DNA elements at distinct times during development.

### *silencer 1* represses expression between rearranged chromosomes

In most cases, transvection is disrupted by chromosomal rearrangements (Lewis, 1954). However, transvection at *ss* occurs despite chromosomal rearrangements (Johnston and Desplan, 2014; Viets *et al*., 2019). We next investigated the mechanisms that facilitate repressive transvection between rearrangements within the *ss* locus, focusing on Ss^ON^/Ss^OFF^ R7 fates. The *ss translocation* allele is a translocation between chromosome X and chromosome 3, with a breakpoint between the *early enhancer* and *silencer 1* (**Fig. 6A**), resulting in *PRE12* and *insulator1* being moved to the X chromosome. Female *translocation/def* flies had 100% Ss^ON^ R7s (**Fig. 6B, S2A**), suggesting loss of repression by the translocated *silencer 1*. Female *translocation/wildtype* and *translocation/prot null* flies displayed ∼73% Ss^ON^ R7s (**Fig. 6C-D, S2A**), suggesting that *silencer1* functions in repressive transvection in the presence of an intact copy of *ss*.

**Figure 6.**
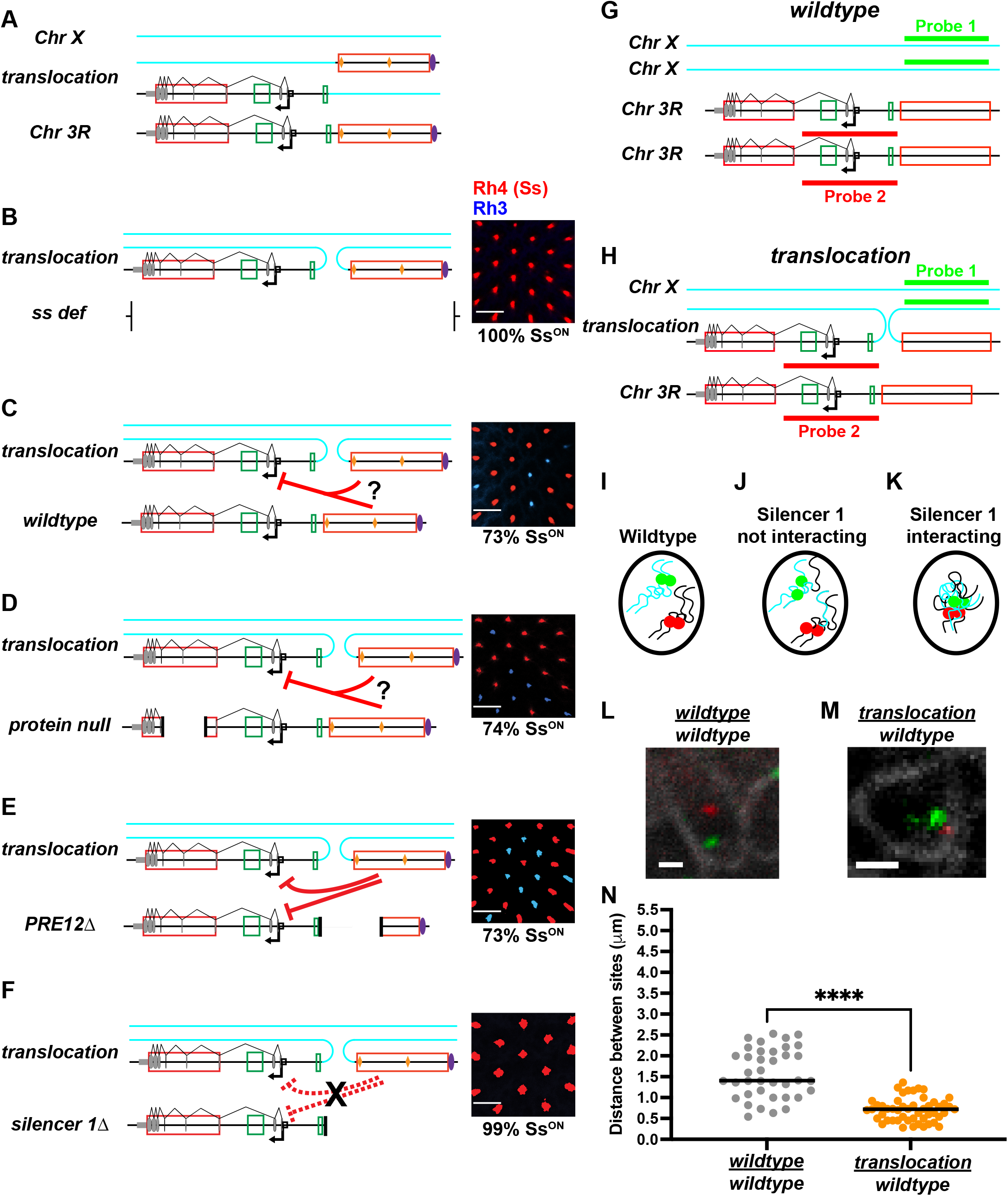
A translocated *silencer 1* performs repressive transvection. **(A)** Schematic of *ss translocation* in female flies. *silencer 1* is translocated to the X chromosome. Schematic includes unaffected X chromosome and unaffected chromosome 3R. Red boxes: *silencers 1* and *2*, green boxes: *early enhancer* and *late enhancer*, gray ovals: exons, black arrow and square: promoter, blue lines: X chromosome, black lines: Chr 3R. **(B-F)** Schematics in predicted chromosomal orientations and representative images of % Ss^ON^ R7s for **(B)** *translocation/ss def*, **(C)** *translocation/wildtype*, **(D)** *translocation/protein null*, **(E)** *translocation/PRE12Δ*, and **(F)** *translocation/silencer1Δ.* Red bars with “?” indicate that it is unclear which copy of *silencer1* is repressing *ss*. Red bars indicate that *silencer1* on the *translocation* is performing repressive transvection. Dotted red bars with black “X” indicate that *silencer1* on the *translocation* is not performing repressive transvection. Scale bars: 15 μm. **(G-H)** Schematics of DNA FISH indicating the locations targeted by DNA probe sets 1 and 2 on a **(G)** *wildtype ss* allele and X chromosome, and **(H)** the *ss translocation*. **(I-K)** Schematics of nuclear localization of probe sets 1 and 2 in **(I)** a *wildtype* control, **(J)** a nucleus where *silencer 1* of *translocation* is not interacting with the *wildtype* allele and **(K)** a nucleus where *silencer 1* translocation is interacting with the wildtype allele. **(L-M)** Representative DNA FISH images of **(L)** *wildtype* control, **(M)** *translocation/wildtype*. Green: probe set 1, red: probe set 2, (**L**) white: prospero (R7 photoreceptor marker) (**M**) white: Lamin B (nuclear marker). Scale bars=1 μm. **(N)** Quantification of **L-M,** n= 2, 3 discs (38, 50 cells). Grey: wildtype control, orange: *translocation / wildtype*. ****=p<0.0001, unpaired, two-tailed t-test.

Similar to females, male *translocation/ss def* flies had 100% Ss^ON^ R7s (**Fig. S2B**). In contrast to females, male *translocation/prot null* flies displayed 100% Ss^ON^ R7s, suggesting that an additional wildtype copy of the X chromosome is required for repression of *ss* between rearranged chromosomes (**Fig. S2A-C**).

Repressive transvection in *translocation/wildtype* and *translocation/prot null* female flies could be driven by *silencer1* on the intact allele or the *translocation* allele (**Fig. 6C-D**). We next tested whether the translocated *silencer 1* represses expression between chromosomes. *translocation/def* flies had 100% Ss^ON^ R7s (**Fig. 6B**) and *PRE12Δ*/*def* flies had 80% Ss^ON^ R7s (**Fig. 5E, 5H**) (Viets *et al*., 2019). *translocation*/*PRE12Δ* female flies, whose only complete copy of *silencer 1* was the translocated copy on the X, displayed 73% Ss^ON^ R7s, a ratio lower than *PRE12Δ*/*def* or *translocation/def* (**Fig. 6E, S2A**). This decrease in Ss^ON^ R7s suggests that the translocated *silencer 1* acts between chromosomes to repress expression from both the *translocation* allele and the *PRE12Δ* allele.

In each case of repressive transvection involving the *translocation* allele, both alleles had an intact *insulator1.* We showed that a copy of *insulator1* on each allele is required for repressive transvection (**Fig. 5**). We next tested if *insulator1* is required for repressive transvection by the *translocation* allele. *silencer 1Δ* lacks *insulator1. silencer 1Δ* homozygotes had 98% Ss^ON^ R7s (**Fig. 5B, 5D**) and *translocation/def* had 100% Ss^ON^ R7s (**Fig. 6B, S2A**). *translocation/silencer 1Δ* flies displayed 99% Ss^ON^ R7s (**Fig. 6F, S2A**), consistent with a requirement of *insulator1* on both alleles for repressive transvection.

To further investigate the mechanisms that facilitate repressive transvection in chromosomal rearrangements, we examined a *ss inversion* allele. This allele contains a chromosomal inversion that moves the *ss* promoter 12 Mb away from *silencer 1* (**Fig. S3A**). *inversion/def* flies displayed 99% Ss^ON^ R7s (**Fig. S3A, S3F**)(Johnston and Desplan, 2014). *inversion*/*wildtype* and *inversion*/*prot null* flies displayed 84% Ss^ON^ R7s and 88% Ss^ON^ R7s, respectively (**Fig. S3B-C, S3F**) (Johnston and Desplan, 2014). *inversion/PRE12Δ* similarly showed a decrease in expression to 85% Ss^ON^ R7s (**Fig. S3D, S3F**), while *inversion/silencer1Δ* showed no repression (99% Ss^ON^)(**Fig. S3E, S3F**). These observations are consistent with *silencer 1* repressing expression at large genomic distances and between chromosomes, dependent on two copies of *insulator1*.

### Chromosomal pairing brings translocated *ss* DNA elements into proximity

The *translocation* and *inversion* alleles break the *ss* gene locus and move *ss* DNA elements to another chromosome or a significant distance away, yet the rearranged *silencer1* represses *ss* via transvection (**Fig. 6, S3**). As physical proximity is required for transvection (Viets *et al*., 2019), we hypothesized that *ss* DNA elements are brought into physical proximity despite these chromosomal rearrangements. To test this hypothesis for the *translocation*, we used DNA FISH (Beliveau et al., 2015; Beliveau et al., 2012) to visualize a region of the X chromosome neighboring the translocated *silencer1* in green and the *ss* gene body in red (**Fig. 6G-H**). We predicted that, in wildtype flies, the X chromosome region and *ss* gene body (Chr 3R) would be far apart in the nucleus (**Fig. 6I**). We predicted that, in female *translocation/wildtype* flies, the X chromosome region and *ss* gene body on chromosome 3 would either be brought into proximity (**Fig. 6K**) or remain at a distance (**Fig. 6J**). The X chromosome region and *ss* gene body were in closer proximity in female *translocation/wildtype* flies than in *wildtype* flies (**Fig. 6L-N**), suggesting that DNA elements in the *ss* gene locus are brought into physical proximity despite a large chromosomal translocation.

We next tested the proximity of DNA elements for the *inversion* allele. We used DNA FISH (Beliveau *et al*., 2015; Beliveau *et al*., 2012) to visualize the region immediately downstream of the proximal inversion breakpoint with green probes and the region immediately upstream of the distal inversion breakpoint, including *silencer1,* with red probes (**Fig. S3G**). We predicted that, in wildtype flies, these regions would be far apart in the nucleus (**Fig. S3H**). We predicted that, in *inversion*/*wildtype* flies, the proximal inversion breakpoint and distal inversion breakpoint would either be brought into proximity (**Fig. S3J**) or remain at a distance (**Fig. S3I**). The proximal inversion breakpoint and distal inversion breakpoint were in closer proximity in flies with the *inversion* allele than in wildtype flies (**Fig. S3K-M**), suggesting that DNA elements in the *ss* gene locus are brought into physical proximity despite the chromosomal inversion.

## Discussion

### Natural variation in Spineless expression is regulated by transvection

Despite the discovery of transvection over 60 years ago (Lewis, 1954), the biological role of transvection has remained elusive. We find that transvection occurs between naturally occurring wild-derived alleles of the *ss* gene. When flies homozygous for a high-frequency *ss* allele are crossed with flies homozygous for a low-frequency *ss* allele, their progeny have an intermediate proportion of Ss^ON^ R7s (**Fig. 2**). Our data suggest a biological role for transvection in preventing extremely high or low Rh4 expression frequencies. The “averaging” of *ss* alleles may keep Ss^ON^ R7:Ss^OFF^ R7 ratios within a certain range required for proper color vision. Supporting this idea, the naturally occurring insertion *sin* causes a decreased proportion of Ss^ON^ R7s and shifts innate color preference from green to blue (Anderson *et al*., 2017), indicating that the proportion of R7 subtypes is important for normal vision. While a broad range of proportions can exist (Anderson *et al*., 2017), transvection averages expression in natural alleles, potentially narrowing the range of cell fate ratios in the wild.

### Developmental timing and cell type specificity of *ss* transvection

Transvection is pervasive throughout the fly genome (Bateman et al., 2012b; Blick *et al*., 2016; Mellert and Truman, 2012; Noble, 2016). Previous studies examined the DNA elements involved in gene activation between chromosomes (Casares *et al*., 1997; Duncan, 2002; Fujioka *et al*., 2016; Gohl *et al*., 2008; Kravchenko *et al*., 2005; Lim *et al*., 2018; Martinez-Laborda *et al*., 1992; Morris *et al*., 1999; Sipos *et al*., 1998) or gene repression between chromosomes (Bantignies *et al*., 2003; Fauvarque and Dura, 1993; Fujioka *et al*., 1999; Gindhart and Kaufman, 1995; Kapoun and Kaufman, 1995; Kassis, 2002; Li *et al*., 2011; Muller *et al*., 1999; Shimell *et al*., 2000; Sigrist and Pirrotta, 1997), but little is known about the mechanisms driving transvection during development.

Using precise CRISPR deletions across the *ss* locus, we identified DNA regulatory elements that govern *ss* transvection. We found that two *ss* enhancers activate *ss* expression between chromosomes (**Fig. 3**), while two PREs and an insulator play roles in repressing *ss* expression between chromosomes (**Fig. 5**). Our findings indicate that activating and repressing transvection involve separable mechanisms.

In the fly eye, many enhancers are able to activate expression between chromosomes, but at different capacities dependent on enhancer strength (Blick et al., 2016). Expression of *ss* during fly eye development provided a unique opportunity to parse out the contributions of different enhancers to transvection in a temporal and cell-type-specific manner. The *early enhancer* activates gene expression between chromosomes in precursors early in development, enabling expression in terminal R7s late in development. The *late enhancer* activates expression late in development via transvection in terminal R7s. These enhancers participate in activating transvection at distinct times in development to promote *ss* expression. The activity of these enhancers between chromosomes occurs at the times and in the cell types in which they are normally active on the same chromosome (i.e. *early enhancer* in precursors early, *late enhancer* in R7s late).

In some situations, the absence of a promoter on the same chromosome increases activation by an enhancer between chromosomes (Casares et al., 1997; Duncan, 2002; Gohl et al., 2008; Martinez-Laborda et al., 1992; Morris et al., 1999; Sipos et al., 1998), possibly due to a reduced competition for the promoter (Bateman *et al*., 2012a; Piwko et al., 2019; Tian *et al*., 2019). In contrast, the *early* and *late enhancers* activated *ss* expression between chromosomes poorly in the absence of the promoter on the same chromosome (**Fig. 4**). There may be two classes of transvection-competent genes: (1) those that increase activating transvection and (2) those that decrease activating transvection, when a promoter is on the same chromosome with an enhancer. As promoter requirements have only been investigated for a subset of transvection-competent genes, including *yellow*, *apterous*, *Abd-b*, and *Ubx* (Casares et al., 1997; Duncan, 2002; Gohl et al., 2008; Martinez-Laborda et al., 1992; Morris et al., 1999; Sipos et al., 1998), it is possible that other genes fall into the *ss* transvection category, and rely on a promoter on the same chromosome. In the case of *ss*, other variables, such as chromatin state, may play a role in transvection efficiency at th*e* locus. For example, in the absence of the *ss* promoter, the gene locus remains in a compacted chromatin state (Voortman, 2021). This may prevent accessibility of the enhancers, hindering their ability to function between chromosomes.

In addition to gene activation, the stochastic pattern of Ss^ON^ and Ss^OFF^ R7s relies on repression of *ss* via two silencer elements (Johnston and Desplan, 2014; Voortman, 2021). In *silencer1,* two PREs and an insulator play a role in repressive transvection. The two PREs facilitate repression, whereas the insulator on both chromosomes is permissively required for repression between chromosomes.

### Transvection despite chromosomal rearrangements within the *ss* gene

While chromosomal rearrangements often disrupt transvection (Duncan, 2002; Lewis, 1954), *ss* transvection occurs in the presence of rearrangements, even when they occur within the gene locus itself (**Fig. 6; S3**). The *ss* locus encompasses a “button” element that promotes chromosome pairing, potentially increasing the robustness of *ss* transvection (Viets *et al*., 2019). Two other “buttons” lie immediately downstream of the *ss* locus and may also promote physical proximity when rearrangements occur within the *ss* locus. Similarly, the *Abd-B* locus overlaps a “button” (Viets *et al*., 2019) and is not disrupted by rearrangements (Gemkow *et al*., 1998; Hendrickson and Sakonju, 1995; Sipos *et al*., 1998).

The ability to “reconstitute” *ss* suggests a genome-wide mechanism that maintains normal gene expression in the presence of chromosomal aberrations. This mechanism likely promotes robustness to rearrangements, with implications for the evolution of long-range gene regulation. In cases where inversions and translocations separate an enhancer from its promoter by large distances, “buttons” might drive a distant enhancer to loop back to its promoter and activate normal gene expression. Over time, this novel enhancer location could evolve to become required for normal expression.

Together, our findings suggest a biological role for transvection and identify spatiotemporal roles for regulatory DNA elements in transvection during development.

## Materials and Methods

### RESOURCE AVAILABILITY

#### Lead Contact

All requests for, or information about, resources can be directed to Robert Johnston Jr.

#### Materials Availability

All fly lines and reagents are available upon request.

### EXPERIMENTAL MODEL AND SUBJECT DETAILS

#### Drosophila Lines

Flies were raised on standard cornmeal-molasses-agar medium and grown at 25° C. All experiments in this study included both male and female flies except where denoted (**Figure 6, S2**).

**Table 1:** Genotypes of *Drosophila* lines.

| Fly line | Full genotype | Source | Figures |
| --- | --- | --- | --- |
| wildtype | *yw*; +; + or *yw*; *sp/CyO*; + | N/A | 1B; 2F-H; 3A-3G; 6C, 6L, 6N; S1B; S2A; S2B, S3K, S3M |
| DGRP Cross 1 Parent 1 | *yw*; *Sp/CyO*; *DGRP57* | Bloomington, (Mackay et al., 2012) | 2A |
| DGRP Cross 1 Parent 2 | *yw*; *Sp/CyO*; *DGRP228* | Bloomington, (Mackay et al., 2012) | 2A |
| DGRP Cross 1 progeny | *yw*; *Sp/CyO*; *DGRP57/DGRP228* | Bloomington, (Mackay et al., 2012) | 2A |
| DGRP Cross 2 Parent 1 | *yw*; *Sp/CyO*; *DGRP57* | Bloomington, (Mackay et al., 2012) | 2A |
| DGRP Cross 2 Parent 2 | *yw*; *Sp/CyO*; *DGRP352* | Bloomington, (Mackay et al., 2012) | 2A |
| DGRP Cross 2 progeny | *yw*; *Sp/CyO*; *DGRP57/DGRP352* | Bloomington, (Mackay et al., 2012) | 2A |
| DGRP Cross 3 Parent 1 | *yw*; *Sp/CyO*; *DGRP338* | Bloomington, (Mackay et al., 2012) | 2A |
| DGRP Cross 3 Parent 2 | *yw*; *Sp/CyO*; *DGRP379* | Bloomington, (Mackay et al., 2012) | 2A |
| DGRP Cross 3 progeny | *yw*; *Sp/CyO*; *DGRP338/DGRP379* | Bloomington, (Mackay et al., 2012) | 2A |
| DGRP Cross 4 Parent 1 | *yw*; *Sp/CyO*; *DGRP379* | Bloomington, (Mackay et al., 2012) | 2A |
| DGRP Cross 4 Parent 2 | *yw*; *Sp/CyO*; *DGRP802* | Bloomington, (Mackay et al., 2012) | 2A |
| DGRP Cross 4 progeny | *yw*; *Sp/CyO*; *DGRP379/DGRP802* | Bloomington, (Mackay et al., 2012) | 2A |
| DGRP Cross 5 Parent 1 | *yw; Sp/CyO; DGRP338* | Bloomington, (Mackay et al., 2012) | 2A |
| DGRP Cross 5 Parent 2 | *yw; Sp/CyO; DGRP802* | Bloomington, (Mackay et al., 2012) | 2A |
| DGRP Cross 5 progeny | *yw; Sp/CyO; DGRP338/DGRP802* | Bloomington, (Mackay et al., 2012) | 2A |
| DGRP Cross 6 Parent 1 | *yw; Sp/CyO; DGRP40* | Bloomington, (Mackay et al., 2012) | 2B |
| DGRP Cross 6 Parent 2 | *yw; Sp/CyO; DGRP228* | Bloomington, (Mackay et al., 2012) | 2B |
| DGRP Cross 6 progeny | *yw; Sp/CyO; DGRP40/DGRP228* | Bloomington, (Mackay et al., 2012) | 2B |
| DGRP Cross 7 Parent 1 | *yw; Sp/CyO; DGRP40* | Bloomington, (Mackay et al., 2012) | 2B |
| DGRP Cross 7 Parent 2 | *yw; Sp/CyO; DGRP352* | Bloomington, (Mackay et al., 2012) | 2B |
| DGRP Cross 7 progeny | *yw; Sp/CyO; DGRP40/DGRP352* | Bloomington, (Mackay et al., 2012) | 2B |
| DGRP Cross 8 Parent 1 | *yw; Sp/CyO; DGRP352* | Bloomington, (Mackay et al., 2012) | 2B |
| DGRP Cross 8 Parent 2 | *yw; Sp/CyO; DGRP379* | Bloomington, (Mackay et al., 2012) | 2B |
| DGRP Cross 8 progeny | *yw; Sp/CyO; DGRP352/DGRP379* | Bloomington, (Mackay et al., 2012) | 2B |
| DGRP Cross 9 Parent 1 | *yw; Sp/CyO; DGRP352* | Bloomington, (Mackay et al., 2012) | 2B |
| DGRP Cross 9 Parent 2 | *yw; Sp/CyO; DGRP338* | Bloomington, (Mackay et al., 2012) | 2B |
| DGRP Cross 9 progeny | *yw; Sp/CyO; DGRP352/DGRP338* | Bloomington, (Mackay et al., 2012) | 2B |
| DGRP Cross 10 Parent 1 | *yw; Sp/CyO; DGRP40* | Bloomington, (Mackay et al., 2012) | 2C |
| DGRP Cross 10 Parent 2 | *yw; Sp/CyO; DGRP379* | Bloomington, (Mackay et al., 2012) | 2C |
| DGRP Cross 10 progeny | *yw; Sp/CyO; DGRP40/DGRP379* | Bloomington, (Mackay et al., 2012) | 2C |
| DGRP Cross 11 Parent 1 | *yw; Sp/CyO; DGRP57* | Bloomington, (Mackay et al., 2012) | 2C |
| DGRP Cross 11 Parent 2 | *yw; Sp/CyO; DGRP379* | Bloomington, (Mackay et al., 2012) | 2C |
| DGRP Cross 11 progeny | *yw; Sp/CyO; DGRP57/DGRP379* | Bloomington, (Mackay et al., 2012) | 2C |
| DGRP Cross 12 Parent 1 | *yw; Sp/CyO; DGRP57* | Bloomington, (Mackay et al., 2012) | 2C |
| DGRP Cross 12 Parent 2 | *yw; Sp/CyO; DGRP802* | Bloomington, (Mackay et al., 2012) | 2C |
| DGRP Cross 12 progeny | *yw; Sp/CyO; DGRP57/DGRP802* | Bloomington, (Mackay et al., 2012) | 2C |
| DGRP Cross 13 Parent 1 | *yw; Sp/CyO; DGRP228* | Bloomington, (Mackay et al., 2012) | 2C |
| DGRP Cross 13 Parent 2 | *yw; Sp/CyO; DGRP802* | Bloomington, (Mackay et al., 2012) | 2C |
| DGRP Cross 13 progeny | *yw; Sp/CyO; DGRP228/DGRP802* | Bloomington, (Mackay et al., 2012) | 2C |
| DGRP Cross 14 Parent 1 | *yw; Sp/CyO; DGRP379* | Bloomington, (Mackay et al., 2012) | 2C |
| DGRP Cross 14 Parent 2 | *yw; Sp/CyO; DGRP195* | Bloomington, (Mackay et al., 2012) | 2C |
| DGRP Cross 14 progeny | *yw; Sp/CyO; DGRP379/DGRP195* | Bloomington, (Mackay et al., 2012) | 2C |
| DGRP Cross 15 Parent 1 | *yw; Sp/CyO; DGRP40* | Bloomington, (Mackay et al., 2012) | 2D |
| DGRP Cross 15 Parent 2 | *yw; Sp/CyO; DGRP195* | Bloomington, (Mackay et al., 2012) | 2D |
| DGRP Cross 15 progeny | *yw; Sp/CyO; DGRP40/DGRP195* | Bloomington, (Mackay et al., 2012) | 2D |
| DGRP Cross 16 Parent 1 | *yw; Sp/CyO; DGRP57* | Bloomington, (Mackay et al., 2012) | 2D |
| DGRP Cross 16 Parent 2 | *yw; Sp/CyO; DGRP338* | Bloomington, (Mackay et al., 2012) | 2D |
| DGRP Cross 16 progeny | *yw; Sp/CyO; DGRP57/DGRP338* | Bloomington, (Mackay et al., 2012) | 2D |
| DGRP Cross 17 Parent 1 | *yw; Sp/CyO; DGRP57* | Bloomington, (Mackay et al., 2012) | 2D |
| DGRP Cross 17 Parent 2 | *yw; Sp/CyO; DGRP195* | Bloomington, (Mackay et al., 2012) | 2D |
| DGRP Cross 17 progeny | *yw; Sp/CyO; DGRP57/DGRP195* | Bloomington, (Mackay et al., 2012) | 2D |
| DGRP Cross 18 Parent 1 | *yw; Sp/CyO; DGRP352* | Bloomington, (Mackay et al., 2012) | 2D |
| DGRP Cross 18 Parent 2 | *yw; Sp/CyO; DGRP195* | Bloomington, (Mackay et al., 2012) | 2D |
| DGRP Cross 18 progeny | *yw; Sp/CyO; DGRP352/DGRP195* | Bloomington, (Mackay et al., 2012) | 2D |
| DGRP Cross 19 Parent 1 | *yw; Sp/CyO; DGRP228* | Bloomington, (Mackay et al., 2012) | 2D |
| DGRP Cross 19 Parent 2 | *yw; Sp/CyO; DGRP338* | Bloomington, (Mackay et al., 2012) | 2D |
| DGRP Cross 19 progeny | *yw; Sp/CyO; DGRP228/DGRP338* | Bloomington, (Mackay et al., 2012) | 2D |
| *ss<sup>sin</sup>/ss<sup>sin</sup>* | *yw; +; ss<sup>sin</sup>/ss<sup>sin</sup>* | (Anderson et al., 2017) | 2E, 2H |
| *ss<sup>sin</sup>/wild type* | *yw; +; ss<sup>sin</sup>/+* | (Anderson et al., 2017) | 2F, H |
| *wild type/ss def* | *yw; +; +/BL7736def* | (Parks et al., 2004) | 3A, 3G; 5D, 5H |
| *early enhancer Δ/ss def* | *yw; +; ss<sup>early enhancer</sup> Δ/BL7736def* | (Parks et al., 2004; Voortman, 2021) | 3B, 3G; 4D |
| *early enhancer Δ/protein null* | *yw; +/(CyO or PM181>>GFP); ss<sup>early enhancer Δ</sup>/ss<sup>D115.7</sup>* | (Duncan, 1998; Lee et al., 2001; Voortman, 2021) | 3C, 3G; 4D |
| *late enhancer Δ/ss def* | *yw; +; ss<sup>late enhancer Δ</sup>/BL7736def* | (Parks et al., 2004; Viets et al., 2019; Voortman, 2021) | 3D, 3G; 4D |
| *late enhancer Δ/protein null* | *yw; +/(Cyo or PM181>>GFP); ss<sup>Late enhancer Δ</sup>/ss<sup>D115.7</sup>* | (Duncan, 1998; Lee et al., 2001; Viets et al., 2019; Voortman, 2021) | 3E, 3G; 4D |
| *early enhancer Δ/late enhancer Δ* | *yw; +; ss<sup>early enhancer Δ</sup>/ss<sup>late enhancer Δ</sup>* | (Viets et al., 2019; Voortman, 2021) | 3F, 3G |
| *promoter Δ/ss def* | *yw; +; ss<sup>promoter Δ</sup>/BL7736def* | (Parks et al., 2004; Voortman, 2021) | 4A |
| *early enhancer Δ/promoter Δ* | *yw; +; ss<sup>early enhancer Δ</sup>/ss<sup>promoter Δ</sup>* | (Voortman, 2021) | 4B, 4D |
| *late enhancer Δ/promoter Δ* | *yw; +; ss<sup>late enhancer Δ</sup>/ss<sup>promoter Δ</sup>* | (Viets et al., 2019; Voortman, 2021) | 4C, 4D |
| *sil1 Δ/sil1 Δ* | *yw; +; ss<sup>RS6279exc44</sup>* | (Thanawala et al., 2013) | 5B, 5D |
| *sil1 Δ/protein null* | *yw; +/(Cyo or PM181>>GFP); ss<sup>RS6279exc44</sup>/ss<sup>D115.7</sup>* | (Duncan, 1998; Lee et al., 2001)( 14) | 5C, 5D |
| *PRE12 Δ/ss def* | *yw; +; ss<sup>PRE12 Δ</sup> / BL7736def* | (Parks et al., 2004; Viets et al., 2019; Voortman, 2021) | 5E, 5H |
| *PRE12 Δ/protein null* | *yw; +/(Cyo or PM181>>GFP); ss<sup>PRE12 Δ</sup>/ss<sup>D115.7</sup>* | (Lee et al., 2001; Viets et al., 2019; Voortman, 2021) (Parks et al., 2004) Bloomington | 5F, 5H |
| *PRE12 Δ/promoter Δ* | *yw; +; ss<sup>PRE12 Δ</sup>/ss<sup>promoter Δ</sup>* | (Viets et al., 2019; Voortman, 2021) | 5G-H |
| *ss<sup>translocation</sup>/ss def* | T(1;3)ss<sup>D114.3</sup>/(+ or y); +;<br>T(1;3)ss<sup>D114.3</sup>/(BL7736,<br>BL7985, or Fullssdel) | Bloomington, (Duncan,<br>1998; Parks et al., 2004;<br>Thanawala et al., 2013) | 6B, S2A-B |
| *ss<sup>translocation</sup>/wild type* | T(1;3)ss<sup>D114.3</sup>/+; +;<br>T(1;3)ss<sup>D114.3</sup>/+ | (Duncan, 1998; Thanawala<br>et al., 2013) | 6C, 6M-N, S2A |
| *ss<sup>translocation</sup>/protein null* | T(1;3)ss<sup>D114.3</sup>/(+ or y); +;<br>T(1;3)ss<sup>D114.3</sup>/ss<sup>D115.7</sup> | (Duncan, 1998; Thanawala<br>et al., 2013) | 6D, S2A-B |
| *ss<sup>translocation</sup>/PRE12Δ* | T(1;3)ss<sup>D114.3</sup>/+; +;<br>ss<sup>PRE12Δ</sup>/T(1;3)ss<sup>D114.3</sup> | (Duncan, 1998; Thanawala<br>et al., 2013; Viets et al.,<br>2019; Voortman, 2021) | 6E, S2A |
| *ss<sup>translocation</sup>/sil1Δ* | T(1;3)ss<sup>D114.3</sup>/+; +;<br>T(1;3)ss<sup>D114.3</sup>/ss<sup>RS6279exc44</sup> | (Duncan, 1998; Thanawala<br>et al., 2013) | 6F, S2A |
| *protein null/ss def* | yw; +/(CyO or PM181>>GFP);<br>ss<sup>D115.7</sup>/BL7736def | Bloomington, (Duncan,<br>1998; Lee et al., 2001;<br>Parks et al., 2004) | S1C |
| *ss<sup>inversion</sup>/ss def* | yw; +; ss<sup>GS2553exc80</sup>/BL7736 | Bloomington, (Parks et al.,<br>2004; Thanawala et al.,<br>2013) | S3A, S3F |
| *ss<sup>inversion</sup>/wild type* | yw; +; ss<sup>GS2553exc80</sup>/+ | (Thanawala et al., 2013) | S3B; S3F,<br>S3L-M |
| *ss<sup>inversion</sup>/protein null* | yw; +; ss<sup>GS2553exc80</sup>/ss<sup>D115.7</sup> | (Duncan, 1998; Thanawala<br>et al., 2013) | S3C, S3F |
| *ss<sup>inversion</sup>/PRE12Δ* | yw; +; ss<sup>GS2553exc80</sup>/ss<sup>PRE12Δ</sup> | (Duncan, 1998; Thanawala<br>et al., 2013) | S3D, S3F |
| *ss<sup>inversion</sup>/sil1Δ* | yw; +; ss<sup>GS2553exc80</sup>/ss<sup>RS6279exc44</sup> | (Duncan, 1998; Thanawala<br>et al., 2013) | S3E, S3F |

### METHOD DETAILS

#### Antibodies

Antibodies were used at the following dilutions: mouse anti-Lamin B (DSHB ADL67.10 and ADL84.12), 1:100; mouse anti-Rh3 (gift from S. Britt, University of Colorado), 1:100; rabbit anti-Rh4 (gift from C. Zuker, Columbia University), 1:100; and Alexa 488 Phalloidin (Invitrogen, Thermo Fisher Scientific, Waltham, MA, USA), 1:80. All secondary antibodies were Alexa Fluor-conjugated (1:400) and made in donkey (Molecular Probes).

#### Antibody Staining

Adult retinas were dissected as described (Hsiao et al., 2012) and fixed for 15 min with 4% formaldehyde at room temperature. Retinas were washed three times in PBS plus 0.3% Triton X-100 (PBX), then incubated with primary antibodies diluted in PBX overnight at room temperature. Next, retinas were washed three times in PBX. Retinas were then incubated with secondary antibodies diluted in PBX overnight at room temperature. Finally, retinas were rinsed three times in PBX. Retinas were mounted in SlowFade Gold Antifade Reagent (Invitrogen). Images were acquired using a Zeiss LSM 700 or Zeiss LSM 980 confocal microscope.

#### Oligopaints probe libraries

**Table 2:**
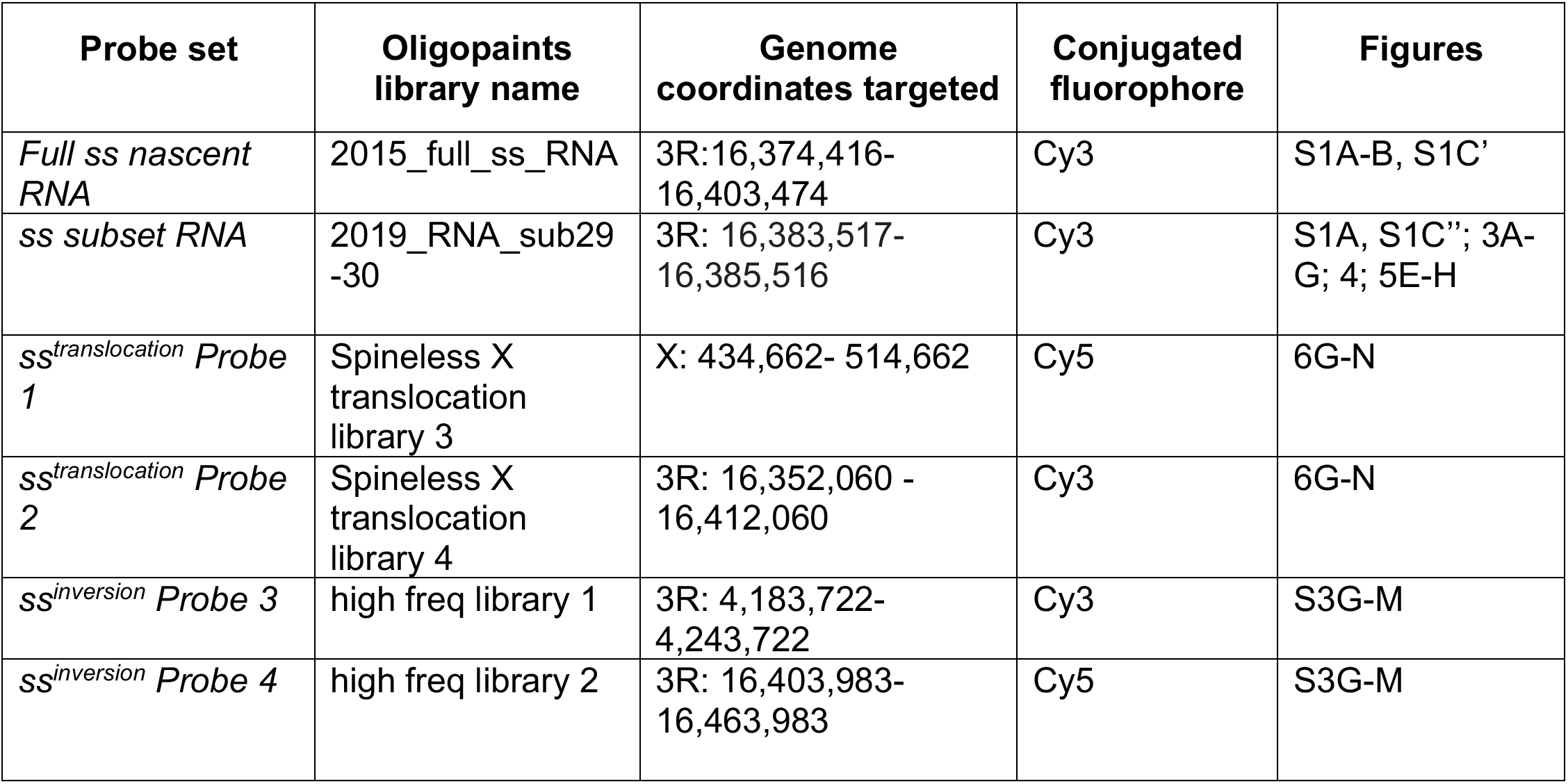
Genome coordinates targeted by Oligopaints probe libraries.

#### Oligopaints probe design

Probes for DNA FISH were designed using the Oligopaints technique (Beliveau et al., 2015; Beliveau et al., 2012), as described in (Viets et al., 2019). Probes for RNA FISH were designed using the Oligominer (Beliveau et al., 2018), as described in (Viets et al., 2019; Voortman, 2021).Target sequences were run through the bioinformatics pipeline available at http://genetics.med.harvard.edu/oligopaints/ to identify sets of 50-bp (DNA FISH, Full *ss* RNA FISH) or 26-bp (subset *ss* RNA FISH) optimized probe sequences (i.e. “libraries”) tiled across the DNA or nascent RNA sequence of interest. Five 19-bp barcoding primers, gene F and R, universal (univ) F and R, and random R or sublibrary F, were appended to the 5’ and 3’ ends of each probe sequence. The gene F and R primers allowed PCR amplification of a probe library of interest out of the total oligo pool, sublibrary F primers (RNA FISH probes only) allowed PCR amplification of subregions of the gene library out of the gene pool, and the univ F and R primers allowed conjugation of fluorophores, generation of single-stranded DNA probes, and PCR addition of secondary sequences to amplify probe signal. The random R primer was appended at the 3’ end of DNA FISH probes to maintain a constant sequence length between all probes that were synthesized in the custom oligo pool.

Barcoding primer sequences were taken from a set of 240,000 randomly generated, orthogonal 25-bp sequences (Xu et al., 2009) and run through a custom script to select 19-bp sequences with ≤15-bp homology to the *Drosophila* genome. Primers were appended to probe sequences using the orderFile.py script available at http://genetics.med.harvard.edu/oligopaints/. Completed probe libraries were synthesized as custom oligo pools by Custom Array, Inc. (Bothell, WA).

#### RNA FISH

RNA FISH was performed as described in (Voortman, 2021). 20 eye-antennal discs attached to mouth hooks from third instar larvae were collected on ice and fixed in 129 μL MilliQ water, 20 μL 10X PBS, 1 μL Tergitol NP-40, 600 μL heptane, and 50 μL fresh 16% formaldehyde. Tubes containing the fixative and eye discs were shaken vigorously by hand, then fixed for 10 minutes at room temperature with nutation. Eye discs were then given four quick washes in 2X SSCT (2XSSC+.001% Tween-20). Eye discs were then removed from the mouth hooks. Next, discs were given one 10-minute wash in 20% formamide+2X SSCT, one 10-minute wash in 40% formamide+2X SSCT, and two 10-minute washes in 50% formamide+2X SSCT. Discs were then prehybridized by incubating for four hours at 37°C. Probes were added in 36 μL hybridization buffer consisting of 50% formamide+2X SSCT+2% dextran sulfate (w/v), + 0.25 μL RNAse inhibitor (Promega N2615). All probes were added at a concentration of 30 pmol fluorophore / μL. 3 μL of probe were used for each genotype. After addition of probes, eye discs were incubated at 37°C for 16 hours with shaking. Eye discs were then washed twice for 30 minutes in 50% formamide+2X SSCT at 37°C with shaking, followed by 10-minute washes at room temperature in 20% formamide+2X SSCT, and 2X SSCT. Discs were DAPI stained for 30 minutes and then washed for 10 minutes 2X SSC with nutation. Discs were mounted in SlowFade Gold (Thermo Fisher S36936) immediately after the final 2X SSC wash, Images were acquired as 63X oil immersion z-stacks (0.15 μm) using a Zeiss LSM980. Images were processed using the airyscan-SR function in Zen and quantified as described below.

#### DNA FISH

DNA FISH was performed as described in (Viets et al., 2019; Voortman, 2021). 20-50 eye-antennal discs attached to mouth hooks from third instar larvae were collected on ice and fixed in 129 μL ultrapure water, 20 μL 10X PBS, 1 μL Tergitol NP-40, 600 μL heptane, and 50 μL fresh 16% formaldehyde. Tubes containing the fixative and eye discs were shaken vigorously by hand, then fixed for 10 minutes at room temperature with nutation. Eye discs were then given three quick washes in 1X PBX, followed by three five-minute washes in PBX at room temperature with nutation. Eye discs were then removed from the mouth hooks and blocked for 1 hour in 1X PBX+1% BSA at room temperature with nutation. They were then incubated in primary antibody diluted in 1X PBX overnight at 4°C with nutation. Next, eye discs were washed three times in 1X PBX for 20 minutes and incubated in secondary antibody diluted in 1X PBX for two hours at room temperature with nutation. Eye discs were then washed two times for 20 minutes in 1X PBX, followed by a 20-minute wash in 1X PBS. Next, discs were given one 10-minute wash in 20% formamide+2X SSCT (2X SSC+.001% Tween-20), one 10- minute wash in 40% formamide+2X SSCT, and two 10-minute washes in 50% formamide+2X SSCT. Discs were then predenatured by incubating for four hours at 37°C, three minutes at 92°C, and 20 minutes at 60°C. Primary probes were added in 45 μL hybridization buffer consisting of 50% formamide+2X SSCT+2% dextran sulfate (w/v), + 1 μL RNAse A. All probes were added at a concentration of ≥5 pmol fluorophore/μL. For FISH experiments in which a single probe was used, 4 μL of probe was added. For FISH experiments in which two probes were used, 2 μL of each probe was added. After addition of probes, eye discs were incubated at 91°C for three minutes and at 37°C for 16-20 hours with shaking. Eye discs were then washed for 1 hour at 37°C with shaking in 50% formamide+2X SSCT. 1 μL of each secondary probe was added at a concentration of 100 pmol/μL in 50 μL of 50% formamide+2X SSCT. Secondary probes were hybridized for 1 hour at 37°C with shaking. Eye discs were then washed twice for 30 minutes in 50% formamide+2X SSCT at 37°C with shaking, followed by three 10-minute washes at room temperature in 20% formamide+2X SSCT, 2X SSCT, and 2X SSC with nutation. Discs were mounted in SlowFade Gold (Thermo Fisher S36936) immediately after the final 2X SSC wash and imaged using a Zeiss LSM700 confocal microscope.

#### Confocal Image Acquisition

Adult retina images were acquired on a Zeiss LSM 700 or Zeiss LSM 980 at 20X magnification in a single z plane. RNA FISH samples were imaged using the Zeiss Airyscan-SR function on the LSM980 at 63X. Airyscan settings were optimally acquired, with a z-thickness of 0.15 μm and zoom 1.7X. z-stack images were acquired starting 1 z-slice below and going to 1 z-slice above precursor R7 cells. Laser power was maintained at 2% with a master gain of 800V and automatic airyscan processing was performed. DNA FISH samples were imaged on a Zeiss LSM 700 at 63X magnification with a Z thickness of 0.2 μm.

### QUANTIFICATION AND STATISTICAL ANALYSIS

#### Quantification of Rh3 and Rh4 expression

Frequency of Rh3 (Ss^OFF^) and Rh4 (Ss^ON^) expression in R7s was scored in adults. Six or more retinas were scored for each genotype (n). 100 or more R7s were scored for each retina. Expression was assessed manually using the count tool in Adobe Photoshop.

#### RNA FISH quantification

All images were max projected starting 1 z-slice below to 1 z-slice above the precursor cells to capture the entire population of precursor R7 cells. Quantifications were performed in 2D on maximum projected images. Quantifications were performed manually using the count tool in Adobe Photoshop. Density measurements were made using a standardized sliding square. A width of 19 μm was chosen for all measurements based off the widest pulse. A 19 μm x 19 μm sliding square was used to count the number of nascent spots/area. Each image contained 3-5 area counts. 3-8 retinas were scored for each genotype.

#### DNA FISH distance quantification

Quantifications were performed as described in (Viets et al., 2019; Voortman, 2021). All quantifications were performed in 3D on z-stacks with a slice thickness of 0.2 μm. Quantifications were performed manually using Fiji (Schindelin et al., 2012; Schneider et al., 2012). To chart the z position of each FISH dot, a line was drawn through the dot and the Plot Profile tool was used to assess the stack in which the dot was brightest. To determine the x-y distance between the two FISH dots, a line was drawn from the center of one dot to the center of the other dot and the length of the line was measured with the Plot Profile tool. The distance between the FISH dots was then calculated in 3D. 50 nuclei from three eye discs were quantified for each genotype (i.e. N=3, n=50), except **Figure 6** wildtype R7 photoreceptors where N=2, n = 38.

#### Statistical analysis for DNA FISH

FISH datasets were tested for a Gaussian distribution using a D’Agostino and Pearson omnibus normality test and a Shapiro-Wilk normality test. These tests indicated a non-Gaussian distribution for the wildtype control and for *ss^inversion^*/WT, so datasets were tested for statistical significance using an unpaired two-tailed T-test. These statistical tests were performed using GraphPad Prism. Statistical tests and p-values are described in the figure legends.

#### Statistical analysis for RNA FISH – early/null pre12/def

FISH datasets were tested for a Gaussian distribution using a Shapiro-Wilk normality test. These tests indicated a Gaussian distribution for almost all genotypes except (1) those lacking expression in precursor cells (*early enhancer Δ*/*def*, *promoter Δ*/*def*), and (2) *earlsy enhancer Δ*/*protein null and PRE12 Δ*/*def*. Since the majority of the data sets were Gaussian distributed, datasets were tested for statistical significance using a one-way ANOVA with Tukey’s multiple comparisons test. These statistical tests were performed using GraphPad Prism. Statistical tests and p-values are described in the figure legends.

#### Statistical analysis for adult % Ss^ON^ R7s

Datasets were tested for a Gaussian distribution using a Shapiro-Wilk normality test. These tests indicated a Gaussian distribution for almost all genotypes except those with near 0 or 100% Rh4/Ss^ON^ (*early enhancer Δ/ def, late enhancer Δ*/ *def*, *silencer1 Δ*/*silencer 1 Δ*, all male *translocation* crosses, *inversion*/ *def*). Since the majority of the data sets were Gaussian distributed, datasets were tested for statistical significance using an unpaired two-tailed t-test (**Figure 2A-D**), or a one-way ANOVA with Tukey’s multiple comparisons test (**Figure 2F; 3; 4; 5; S2**). These statistical tests were performed using GraphPad Prism. Statistical tests and p-values are described in the figure legends.

## Figure Legends

**Supplemental Figure 1 (Related to Figure 3).**
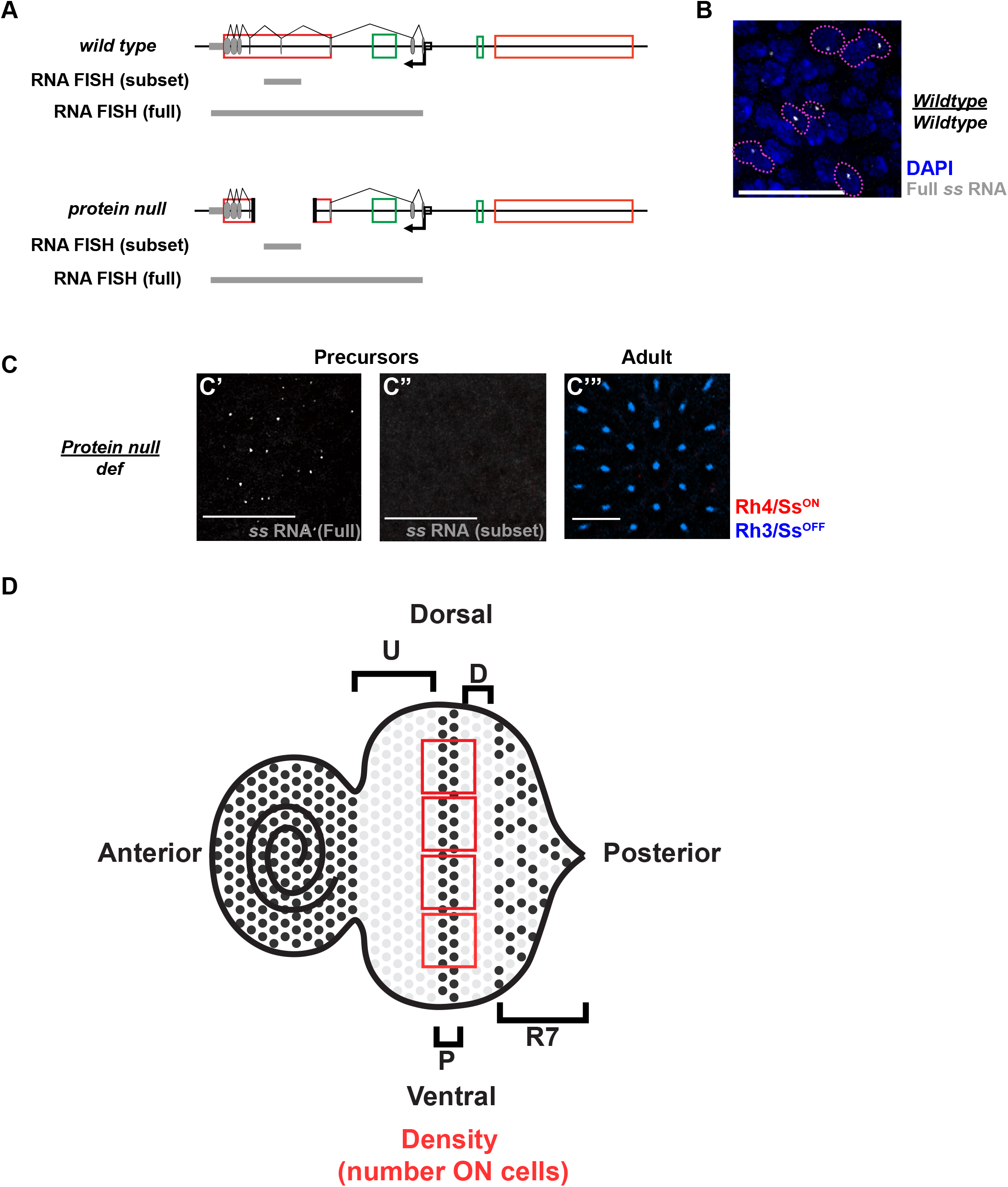
RNA FISH strategy to monitor transvection. **(A)** RNA FISH probe design. Grey bars denote probe target. “Full” includes 219 single stranded DNA probes tiling from the *ss promoter* through the 3’UTR. “Subset” contains 30 probes tiling a 2 kb region within the deleted region of the *protein null* allele (exon 4 + partial intron 3 + partial intron 4). Subset probes detect transcription from intact alleles but not the *protein null* allele. **(B)** Representative image of RNA FISH in *wildtype* precursor cells. Probes target the full nascent transcript and result in a single nascent spot per nucleus due to chromosome pairing. Grey: *ss RNA,* blue: DAPI, Cells expressing *ss* RNA outlined in pink. Scale bar: 15 μm. **(C-C’”)** RNA FISH against the *protein null/def* using **(C’)** full and **(C”)** subset probes. Full probes detect transcripts from the *protein null* allele in precursor cells. Subset probes do not detect transcripts from the *protein null* allele in precursor cells. (C’’’) Ss protein is not expressed in the *protein null/def* flies. Antibody staining in adult eyes against Rh4 (Ss): red, Rh3: blue. Scale bars: 15 μm. **(D)** Quantification of RNA FISH in precursor cells. Precursor density (number of *ss*^ON^ cells) was counted by taking a sliding box (3 to 5 per disc) along the dorsal/ventral axis of the disc in line with precursor cells and counting the number of nascent transcription sites.

**Supplemental Figure 2 (Related to Figure 6).**
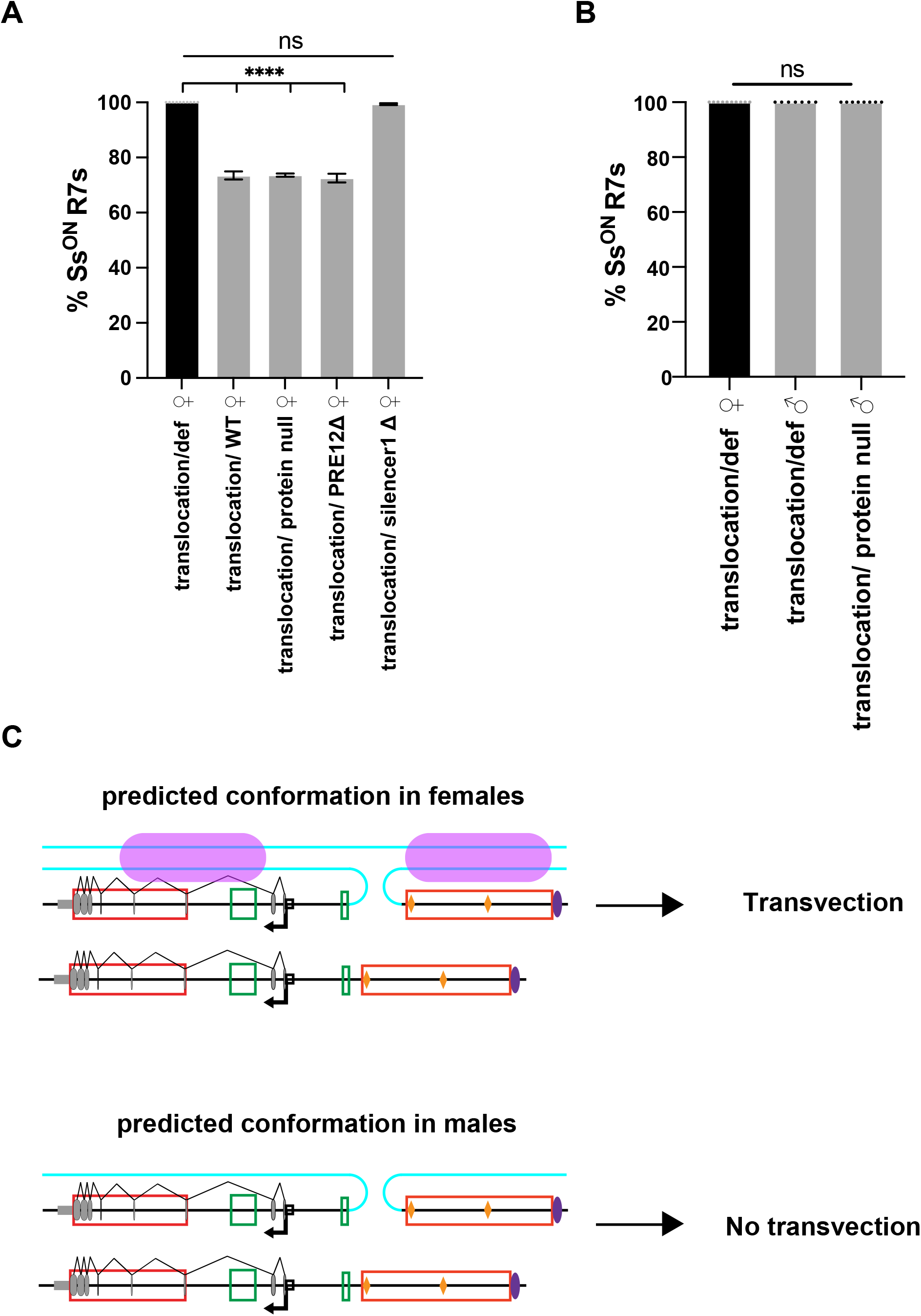
A translocated *silencer1* requires an intact X chromosome for repressive transvection. **(A)** Quantification of % Ss^ON^ R7s in female flies containing the *translocation* allele (as in Figure 6B-F). P values calculated by one-way ANOVA with Tukey’s multiple comparison test. ****=p<0.0001, ***=p<0.001, **=p<0.005, *=p<0.05. Error bars: SEM. **(B)** Quantification of % Ss^ON^ R7s in male and female flies containing the *translocation* allele. P values calculated by one-way ANOVA with Tukey’s multiple comparison test. ****=p<0.0001, ***=p<0.001, **=p<0.005, *=p<0.05. Error bars: SEM. **(C)** Predicted chromosomal conformation in male and female translocation flies. Transvection occurs in the presence of a second copy of the X chromosome. Pink ovals denote potential pairing interactions that facilitate transvection in the presence of a second X chromosome in females but not males.

**Supplemental Figure 3 (Related to Figure 6):**
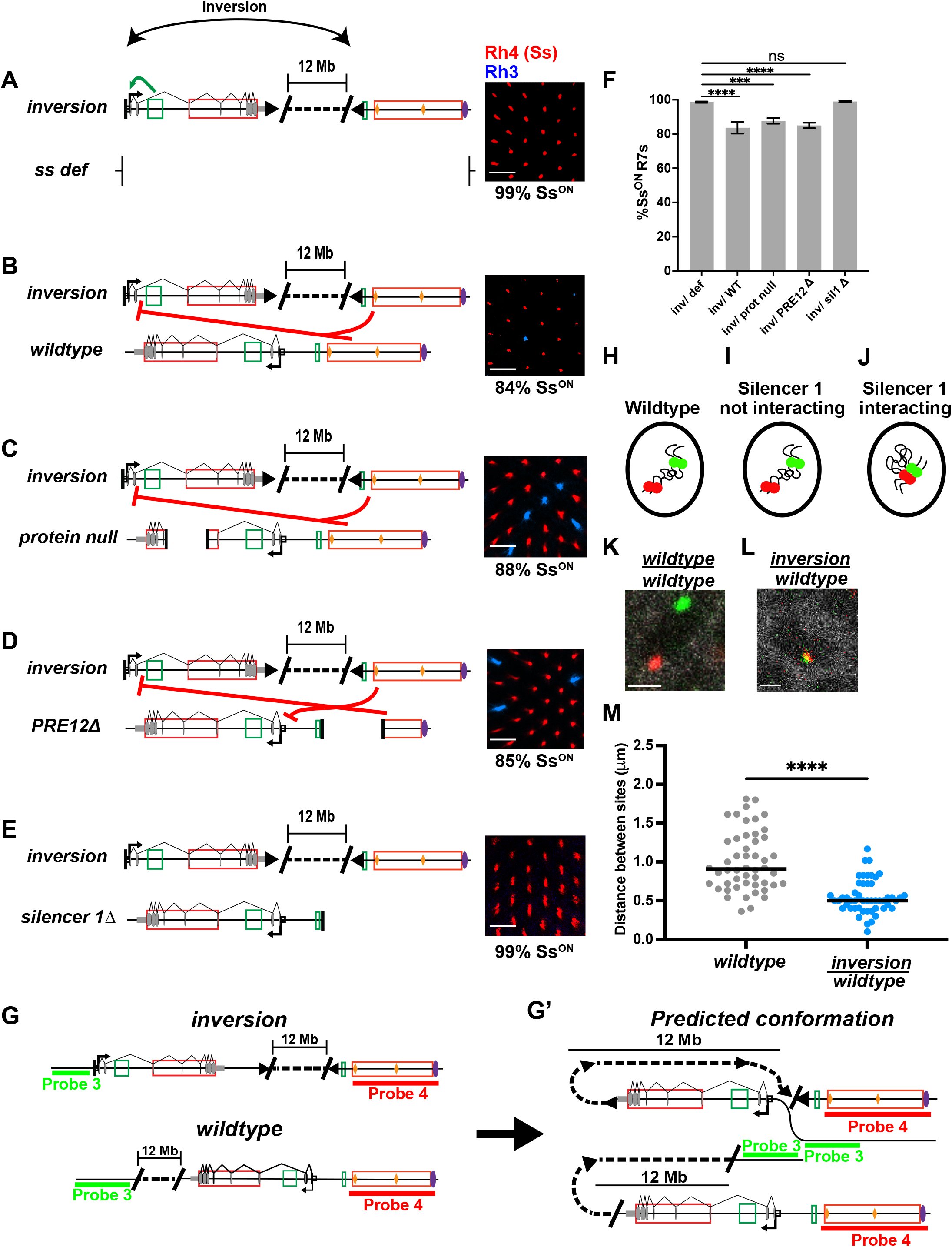
*silencer 1* on a chromosomal inversion performs repressive transvection. **(A-E)** Schematics and representative images of % Ss^ON^ R7s for **(A)** *inversion/ss def*, **(B)** *inversion/wildtype*, **(C)** *inversion/protein null*, **(D)** *inversion/PRE12Δ*, and **(E)** *inversion/silencer1Δ.* Scalebar = 15 μm. **(F)** Quantification of % Ss^ON^ R7s. P values calculated by one-way ANOVA with Tukey’s multiple comparison test. ****=p<0.0001, ***=p<0.001, **=p<0.005, *=p<0.05. Error bars: SEM. **(G-G’).** Schematics of DNA FISH indicating the locations targeted by DNA probe sets 3 and 4 on **(G)** a *wildtype ss* allele and the *ss inversion* allele. Dotted lines indicate the 12-Mb region that separates *silencer 1* and the *ss* protein coding region on the *inversion* allele. **(G’)** Right: predicted locus conformation for repressive transvection. **(H-J)** Schematics of nuclear localization of probe sets 3 and 4 in **(H)** a *wildtype* control, **(I)** a nucleus where *silencer1* on the *inversion* allele is not interacting with the transcribed region of the *ss* locus, and **(J)** a nucleus where *silencer 1* on the *inversion* allele is interacting with the transcribed region of the *ss* locus. **(K-L)** Representative DNA FISH images of **(K)** *wildtype* control, n= 3 discs (50 cells), and **(L)** *inversion/wildtype*, n= 4 discs (50 cells). Green: probe set 3, red: probe set 4, white: Lamin B (nuclear marker). Scale bars=1 μm. **(M)** Quantification of **K-L**. Grey: control, blue: inversion. ****=p<0.0001, unpaired, two tailed T-test.

